# Sport-Related Concussion in Adolescent Athletes is Associated with Acute White Matter Alterations, Delayed Network Changes, and Individual-Level Injury Patterns

**DOI:** 10.64898/2026.08.05.743045

**Authors:** Elizabeth V Castro, Mohammad N Haider, Ferdinand Schweser, John J Leddy, Jeffrey C Miecznikowski, Sarah F. Muldoon

**Affiliations:** Neuroscience Program, University at Buffalo, SUNY, Buffalo, New York; UBMD Orthopaedics and Sports Medicine, Jacobs School of Medicine and Biomedical Sciences, University at Buffalo, SUNY, Buffalo, New York; Buffalo Neuroimaging Analysis Center, Department of Neurology, Jacobs School of Medicine and Biomedical Sciences, University at Buffalo, SUNY, Buffalo, New York; Center for Biomedical Imaging, University at Buffalo, SUNY, Buffalo, New York; Department of Biostatistics, School of Public Health and Health Professions, University at Buffalo, SUNY, Buffalo, New York; Department of AI and Society, University at Buffalo, SUNY, Buffalo, New York; Institute for Artificial Intelligence and Data Science, SUNY, Buffalo, New York

**Keywords:** Diffusion magnetic resonance imaging, differential tractography, concussion, adolescents, variability

## Abstract

Sport-related concussions (SRC) are heterogenous injuries that produce a variety of symptoms and recovery trajectories. This heterogenous nature and focus on group-level analyses in current literature may obscure individual level results that could better inform clinical SRC management. In a prospective case-control study, we used diffusion magnetic resonance imaging (dMRI) to quantify longitudinal, whole-brain microstructural white matter changes reflecting axonal injury and inflammation and structural network-level alterations following SRC in adolescent athletes. Differential tractography assessed individual white matter track changes from acute injury to clinical recovery on an individual level. Acutely after SRC, but not after recovery, concussed adolescents demonstrated increased (i) quantitative anisotropy, (ii) restricted diffusion imaging, and (iii) isotropy, indicating increased microstructural disruptions early after injury. At the network level, differences were seen not acutely but after clinical recovery: whole brain network structure was more similar with reduced capacity for information spread among the concussed adolescents compared with controls. At the individual level, consistent patterns of damaged white matter tracks persisted in the concussed males but not in the concussed females. These results indicate that adolescent athlete brains are impacted acutely at the microstructural level following SRC, but that macroscale network disruptions appear after microstructure damage resolution, and they can persist beyond clinical recovery. Sex differences in the brain’s microstructural response to SRC, highlight the need for future research to include individualized and sex-stratified analyses to guide targeted SRC management.

## 1. INTRODUCTION

Concussion, a form of mild traumatic brain injury (mTBI), has been reported by approximately 19.5% of US adolescents aged 12 to 18,^1^ most often from sport-related activities.^2^ Sport-related concussion (SRC) is characterized by a neurometabolic cascade^3^ that includes axonal injury, neuroinflammation, and altered cerebral blood flow (CBF) regulation.^4^ The symptoms of SRC are heterogenous including, but not limited to, headaches, dizziness, nausea, oculomotor issues, and cognitive deficits.^3,5^ Importantly, on routine clinical neuroimaging scans such as computed tomography and T1-weighted magnetic resonance imaging (MRI), SRC typically shows no structural abnormalities.^3^

However, advanced neuroimaging has shown that SRC is associated with white matter abnormalities, particularly early after injury.^6^ Diffusion MRI (dMRI) can quantify microstructural white matter changes following SRC using a variety of metrics.^7^ Axonal damage from the neurometabolic cascade also alters structural brain networks, which can be quantified using graph theory-based network measures.^8–10^ Diffusion and network measures have been extensively studied in concussion and mTBI, but the results vary with no clear consensus for the direction of change for many of them.^11–15^

The inconsistent imaging findings may be due to different methodologies, different sample characteristics, and a lack of subject-level analyses that encompass the complexity of SRC. Some studies, for example, focus on whole brain or global-level quantification of microstructure and structural connectivity^16–18^ while others investigate specific regions of the brain.^19–21^ One of the regions largely neglected in diffusion imaging after SRC is the brainstem.^22–25^ There are also differences in the inclusion criteria, reporting, and investigation of sample characteristics across studies. Sex differences have been noted in SRC, specifically in symptoms, recovery, and brain structure/function.^26^ Despite this, one review paper reported that only 3 out of 26 studies investigated diffusion measure-related sex differences in SRC.^15^ Finally, the majority of dMRI SRC studies used group-level analyses with a paucity of subject-level investigations.^15^ Group-level analyses of SRC may obscure individual subtle changes after SRC.

In this study we compared whole brain diffusion and network measures in concussed and control adolescent athletes and investigated individual variability in white matter tracks after SRC. We found that acutely after SRC, but not after recovery, concussed adolescents had increased diffusion metrics, but, at the network level, any differences could only be detected after recovery. At this delayed time point, whole brain network structure was more homogenous and demonstrated reduced capacity for information spread among the concussed adolescents compared to the controls. At the subject level, consistent patterns of damaged white matter tracks were seen in the concussed group, and when stratified by sex, these patterns persisted in the concussed males but not in the concussed females. Our results suggest that adolescent brains are impacted acutely at the microstructural level following SRC, but that macroscale network disruptions may appear only after microstructure damage resolution, and that they persist after clinical recovery.

## 2. METHODS

### 2.1 Study Design

This prospective case control study was approved by the University at Buffalo’s Institutional Review Board (IRB). Adolescents diagnosed with an acute SRC at three university-affiliated sports medicine clinics in Buffalo, NY were asked to participate in the study and written informed consent was obtained. Parental consent and participant assent were obtained for all minors (aged 13-17). Participants came for a total of 4 visits at two timepoints, one MRI visit and one physiology lab visit within 10 days of SRC (baseline) and again after clinical recovery as determined by the study physician (follow-up).

Controls were recruited from local high schools and sports teams. Mean recovery time for adolescents with SRC is typically under 1-month,^27^ so controls were invited to return to perform repeat MRI and physiological testing 4-6 weeks after (follow-up) the first visit (baseline).

### 2.2 Participants

Inclusion criteria for the concussed group were: 1) aged 13-18 years; 2) any race, ethnicity, or sex; 3) sustained SRC within 10 days prior to initial research visit; 4) demonstrated exercise intolerance on the Buffalo Concussion Treadmill Test (BCTT); and 5) report more than 7 points on the Post-Concussion Symptom Scale (PCSS, max = 126).^28^ The control group inclusion criteria included: 1) aged 13-18 years; 2) any race, ethnicity, or sex; 3) no history of concussion within the past year; and 4) played at least one organized sport. Both concussed and control participants were excluded if they had any one of the following: 1) had a history of or current diagnosis of moderate or severe TBI; 2) inability to exercise; 3) active substance abuse/dependence; 4) no more than 3 previous concussions; 5) currently on beta- or calcium-blocker or medications that affect the autonomic nervous system (e.g. ADHD stimulants and mood-stabilizers); or 6) were not eligible for MRI.

### 2.3 MRI acquisition

At the baseline and follow-up assessments, all participants underwent MRI examination using a Canon Vantage Titan 3T scanner (Canon Medical Systems, Otawara, Japan) with a 32-channel brain coil. The sequences consisted of high-resolution 3D T1-weighted (T1w) MPRAGE imaging and multi-shell, axial, 3D single-shot spin-echo echo-planar diffusion-weighted imaging (DWI). The T1w images were acquired with 1 mm^3^ isotropic voxels, 256 x 256 mm^2^ field of view (FOV), 192 sagittal slices, echo time (TE)/repetition time (TR)/inversion time (TI) = 3.2ms/6.2ms/900ms, flip angle of 8 degrees and BW = 488Hz/Px. DWI images were acquired with 2.67 x 2.67 x 2.70 mm^3^ voxels, 53 axial slices without gap, 256 x 256 mm^2^ FOV, TE/TR =100 ms/11819 ms. The DWI acquisition included 8 b = 0 s/mm^2^, 21diffusion-weighted directions at b=1000 s/mm^2^, and 25 diffusion-weighted directions at 2,000 s/mm^2^. An additional b = 0 s/mm^2^ volume was acquired with reversed phase-encoding for susceptibility-induced geometric correction.

### 2.4 MRI pre-processing

Visual inspection of all MRI data was performed to ensure data quality. DWI data was preprocessed using FMRIB Software Library (FSL) Version 6.0.5.1.^29^ Susceptibility distortions, eddy currents, and head motion artifacts were corrected for using TOPUP^30^ and EDDY^31^ analysis tools. Quality control following artifact correction was performed using EDDY quality control tools.^32^

### 2.5 Diffusion Tensor Imaging (DTI) and Generalized Q-Sample Imaging (GQI) Measure Analysis

A population-specific atlas^33^, the Purdue Neurotrauma Group Adolescent Collision-Sport Athletes Brain Atlas, was chosen and registered to each subject’s DWI data using Advanced Normalization Tools software Version 2.5.1.^34^ The atlas was developed using 13-19-year-old collision-sport athletes T1-weighted and DWI scans and employed a similar labeling protocol as the Desikan Killiany atlas with 181 regions, including the brainstem and cerebellum.^33,35^ After registration, DSI Studio Version Chen was used to reconstruct the DWI data in native space using generalized q-sample imaging (GQI).^36^ Four diffusion tensor imaging (DTI)^37^ and three GQI measures were calculated. The DTI measures included fractional anisotropy (FA), mean diffusivity (MD), axial diffusivity (AD), and radial diffusivity (RD), while the GQI measures included quantitative anisotropy (QA)^38^, restricted diffusion imaging (RDI)^39^, and isotropy (ISO)^36^. FA is thought to relate to axonal integrity, MD is associated with edema, AD may represent axonal damage, and RD is a marker of myelination.^37^ QA is the GQI counterpart of FA and represents axonal density, RDI combines AD and RD and is thought to be associated with neuroinflammation, and ISO is the counterpart of MD and is also associated with edema.^36,39,40^

### 2.6 Network Measure Analysis

Following GQI reconstruction in DSI Studio, whole brain tractography was performed using a deterministic fiber tracking algorithm^40^ with the following parameters: threshold index = QA, minimum streamline length = 30 mm, maximum streamline length = 200 mm, and 1,000,000 seeds. In addition to these fixed parameters, anisotropy threshold, angular threshold, and step size were randomly generated based on DSI Studio criterion. Tractography informed end-to-end connectivity matrices were calculated with 100 repetitions for each subject and exported into MATLAB Version 23.2.0 (R2023b) (The MathWorks Inc., Natick, MA). Matrices were then averaged for each subject, normalized for atlas brain region size, and the Brain Connectivity Toolbox^41^ and previously published scripts were used to calculate global network measures. The network measures in this study included global clustering coefficient, global efficiency, small-world propensity^42^, synchronizability^43^, spectral radius^44^, and global dissimilarity^45^. The global clustering coefficient reflects the local connectivity around nodes in a network, the global efficiency is indicative of how well nodes in a network communicate, small-world propensity is the balance between integration and segregation in a network, synchronizability represents how easily a network can engage in coordinated activity, the spectral radius indicates how activity spreads in a network, and global dissimilarity informs on how dissimilar the structure of networks in a group are from each other.

### 2.7 Differential Tractography

In this study, differential tractography^38^ was implemented in DSI Studio to compare baseline and follow-up dMRI data on an individual level to determine longitudinal white matter changes. White matter increases (follow-up minus baseline) and decreases (baseline minus follow-up) were calculated for every participant for 3 GQI measures (QA, RDI, and ISO) with thresholds ranging from 5% to 40%, with a step size of 5%. Tractography was performed using the same parameters as for the network measures. Since a higher relative threshold filters out false positives, and a threshold that is too high may result in false negatives, the thresholds were tested and chosen using visual inspection of the results and the false discovery rate (FDR) method outlined by Yeh et al.^38^ Here we analyzed three thresholds (25%, 30%, and 35%) to investigate different levels of structural change. Differential tractography results for all participants who had a difference between baseline and follow-up at each threshold were visually inspected. At each chosen threshold and for each group, the resulting white matter tracks were named using DSI Studio’s “Recognize and Rename” feature and aggregated by counting the frequency for each white matter track that appeared as being different among participants. The aggregated control tracks were used as a noise-derived baseline to compare the concussed tracks against. Registration failed for two concussed participants, one male and one female, therefore they were excluded from further analysis. Only white matter decreases are presented below as the white matter increases had minimal observable findings at the 25%, 30%, or 35% thresholds. FDR values and tracks for the white matter increases are presented in Tables S21 and S22.

### 2.8 Statistical Analysis

Demographics were compared between the concussed and control groups. Continuous variables were compared using a two-sample t-test and discrete variables were compared using a chi-squared test for significance. For the diffusion and network measures, it was determined that the data were not normally distributed, and therefore non-parametric statistics were used to analyze all measures. Group differences between the concussed and control groups for each diffusion and network measure were analyzed at baseline, follow-up, and as differences (follow-up subtracted from baseline) using Wilcoxon Rank Sum tests, and Wilcoxon Signed Rank tests for the difference with significance level of 0.05 for all tests. The rank-biserial correlation effect size with 95% confidence interval was computed for Wilcoxon Rank Sum tests and the matched-pairs rank-biserial correlation effect size with 95% confidence interval for Wilcoxon Signed Rank tests. Violin plots were made for baseline and follow-up results, and paired line plots for the difference. Graphs were made using GraphPad Prism Version 10.0.0 (GraphPad Software, Boston, MA). Analyses were performed in R Statistical Software Version 4.4.1.^46^ Sex was not included in the primary diffusion and network analyses as it was not identified as a confounder

or effect modifier; however, differential tractography results were additionally stratified by sex as a secondary analysis. For differential tractography, false discovery rates (FDR) were calculated as the average number of tracks for the control group divided by the average number of tracks for the concussed group. FDR rates (thresholds) were selected if they were ≤ 0.20 and based on visual inspection of tractography results.

## 3. RESULTS

Table 1 presents the groupwise demographics for this study at baseline. We had 26 concussed athletes with complete data (both baseline and follow-up data) and 24 controls with complete data. At baseline there were 31 concussed athletes and 32 controls and at follow-up there were 29 concussed athletes and 26 controls. At baseline, participants with a concussion reported significantly more previous concussions compared to controls.

**Table 1.** Participant Demographics at Baseline. Continuous variables are presented as means and standard deviations. Categorical variables are presented as frequencies and percentages.

|  | Concussed Group | Control Group | p-value |
| --- | --- | --- | --- |
| n | 31 | 32 | - |
| Age (year) | 15.42 ± 1.18 | 15.72 ± 1.42 | 0.367 |
| Sex, n (%) |  |  |  |
| Female | 15 (48.4%) | 15 (46.8%) | 0.904 |
| Male | 16 (51.6%) | 17 (53.1%) |  |
| Previous Concussion, n (%) |  |  |  |
| 0 | 9 (29.0%) | 28 (87.5%) | <0.001 |
| 1 or more | 22 (71.0%) | 4 (12.5%) |  |
| Height (meters) | 1.67 ± 0.11 | 1.69 ± 0.08 | 0.375 |
| Weight (kilograms) | 68.08 ± 12.06 | 67.36 ± 12.70 | 0.820 |
| Difference from Visit 1 to Visit 2 (days) | 40.14 ± 33.30 | 48.37 ± 17.18 | 0.270 |

To assess microstructural changes after SRC, we reconstructed our data using two complimentary methods: (i) a model-based approach with more clinically interpretable measures, DTI, and (ii) a model-free approach which is better at resolving crossing fibers, GQI. First, we examined measures from both reconstruction methods at baseline, acutely after SRC for the concussed group, and at enrollment for controls (Figure 1, DTI [A-D] and GQI [E-G]). There were no significant differences between the concussed and control groups for the DTI measures FA, MD, RD, or AD (Fig. 1A-D, Table S1). However, there were significant differences between groups for all three GQI measures, with the concussed group having significantly higher QA, ISO, and RDI compared to the control group, indicating that there was microstructural damage after SRC (Fig. 1E-G, Table S1).

**Figure 1.**
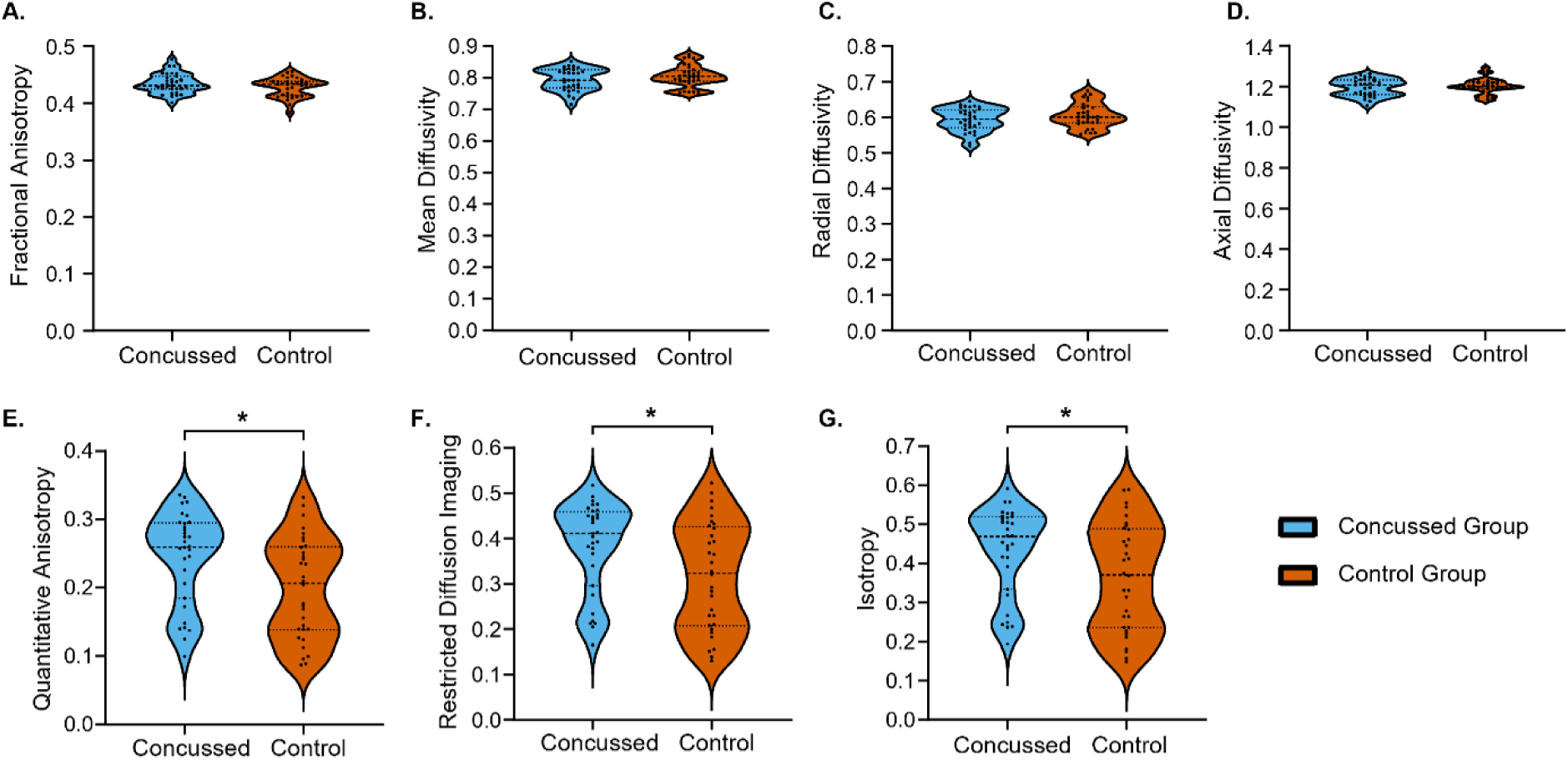
DTI and GQI Measures Compared Between Groups at Baseline. Violin plots showing the distributions of the DTI (A-D) and GQI (E-G) measures by group. (A) Fractional Anisotropy (Concussed median:0.431, Control median:0.433, p-value=0.277). (B) Mean Diffusivity (Concussed median:0.792, Control median:0.804, p-value=0.432). (C) Radial Diffusivity (Concussed median:0.595, Control median:0.601, p-value=0.265). (D) Axial Diffusivity (Concussed median:1.208, Control median:1.201, p-value=0.811). (E) Quantitative Anisotropy (Concussed median:0.260, Control median:0.206, p-value=0.011). (F) Restricted Diffusion Imaging (Concussed median:0.411, Control median:0.323, p-value=0.026). (G) Isotropy (Concussed median:0.469, Control median:0.371, p-value=0.342).

DTI and GQI measures were also assessed at follow-up, after clinical recovery for the concussed group, and after 4-6 weeks for the control group (Figure 2, DTI [A-D] and GQI [E-G]). After recovery, no significant differences between groups were observed for either DTI (FA, MD, RD, AD) or GQI (QA, ISO, or RDI) measures (Table S1). This suggests that the microstructural damage seen in the GQI measures at baseline is no longer present at clinical recovery.

**Figure 2.**
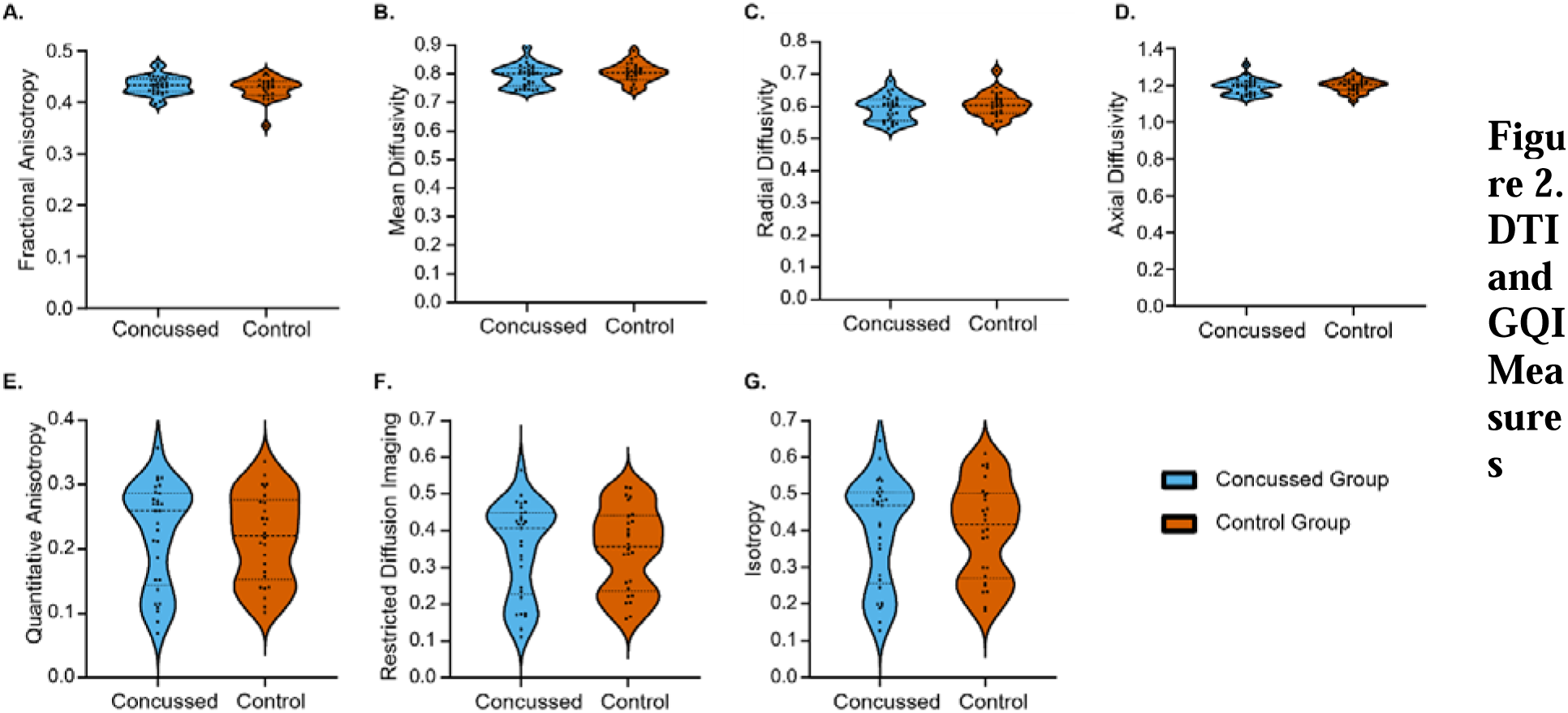
DTI and GQI Measures Compared Between Groups at Follow-Up. Violin plots showing the distributions of the DTI (A-D) and GQI (E-G) measures by group. (A) Fractional Anisotropy (Concussed median:0.434, Control median:0.431, p-value=0.327). (B) Mean Diffusivity (Concussed median:0.802, Control median:0.805, p-value=0.370). (C) Radial Diffusivity (Concussed median:0.601, Control median:0.603, p-value=0.319). (D) Axial Diffusivity (Concussed median:1.20, Control median:1.21, p-value=0.352). (E) Quantitative Anisotropy (Concussed median:0.259, Control median:0.220, p-value=0.757). (F) Restricted Diffusion Imaging (Concussed median:0.407, Control median:0.356, p-value=0.967). (G) Isotropy (Concussed median:0.468, Control median:0.416, p-value=0.967).

We then compared the DTI and GQI measures from baseline to follow-up within groups to assess how white matter damage evolves from injury to clinical recovery (Figure 3, DTI [A-D] and GQI [E-G]). There were no significant differences observed between baseline and follow-up for the controls for the DTI or GQI measures (Table S2). Interestingly, the change in the DTI and GQI measures from baseline to follow-up in the concussed group did not reach statistical significance (*p-*values = 0.055-0.066, Table S2). The lack of significant change from injury to recovery conflicts with our observation of significant group differences in GQI measures at baseline. However, as seen in Fig. 3, the paired line plots for the DTI measures did not change much whereas the GQI measures showed substantial individual variability between subjects from baseline to follow-up. This observation indicates that injury recovery is heterogenous and future work will be necessary to explore the root of this variability.

**Figure 3.**
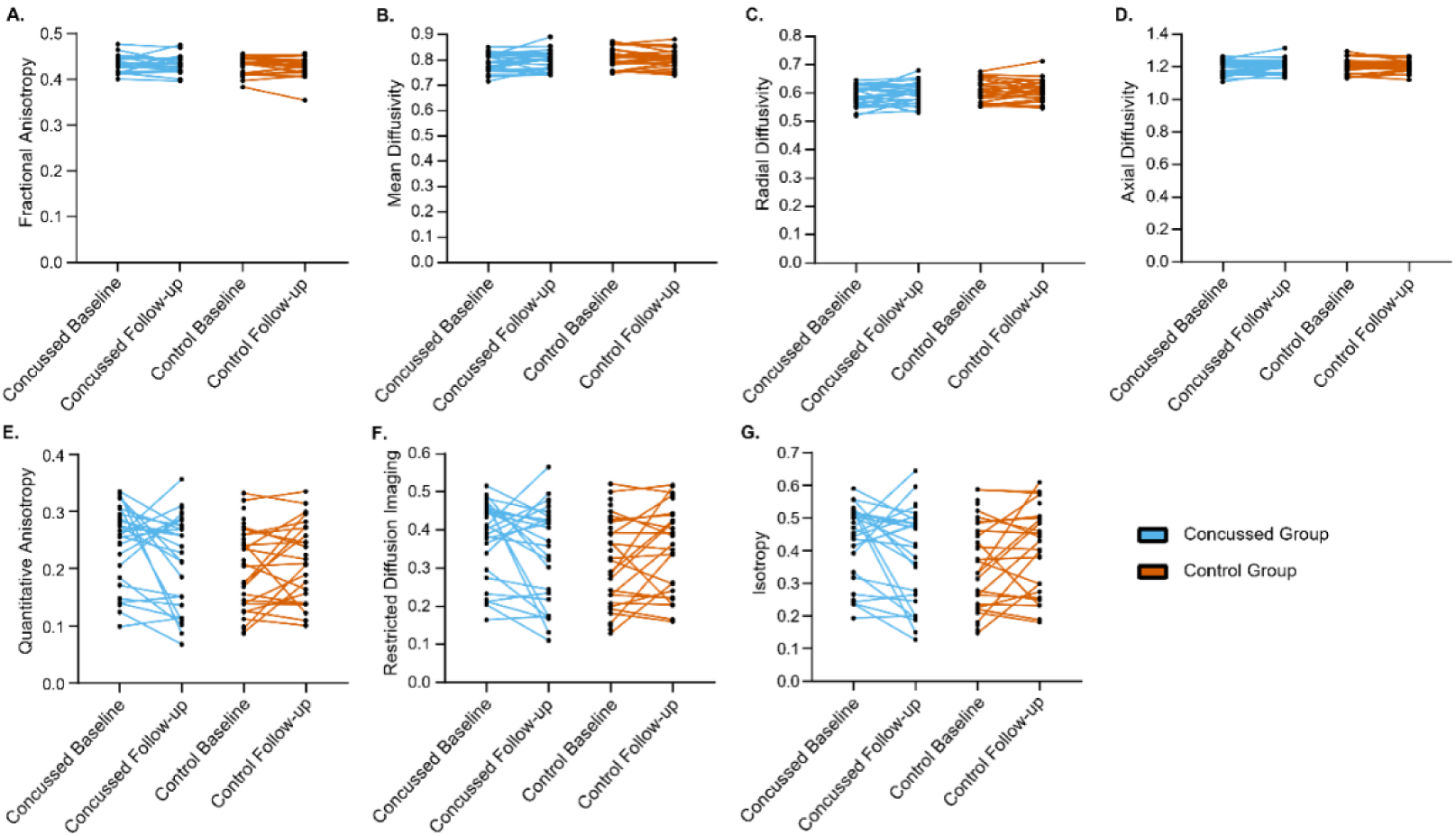
DTI Measures Compared Between Groups for the Difference in Timepoints. Paired line plots displaying the change from baseline to follow-up for the diffusion measures. (A-D) The results for the 4 DTI measures. (E-G) The results for the 3 GQI measures. (A) Fractional Anisotropy (Concussed baseline median:0.431, Concussed follow-up median:0.434, p-value=0.117; Control baseline median:0.0.433, Control median: 0.431, p-value=0.584). (B) Mean Diffusivity (Concussed baseline median:0.792, Concussed follow-up median:0.802, p-value=0.303; Control baseline median:0.804, Control follow-up median:0.805, p-value=0.855). (C) Radial Diffusivity (Concussed baseline median:0.595, Concussed follow-up median:0.601, p-value=0.157; Control baseline median:0.601, Control follow-up median:0.603, p-value=0.790). (D) Axial Diffusivity (Concussed baseline median:1.208, Concussed follow-up median:1.200, p-value=0.784; Control baseline median:1.202, Control follow-up median:1.206, p-value=0.999). (E) Quantitative Anisotropy (Concussed baseline median: 0.260, Concussed follow-up median:0.259, p-value=0.056; Control baseline median:0.206, Control follow-up median:0.220, p-value=0.303). (F) Restricted Diffusion Imaging (Concussed baseline median:0.411, Concussed follow-up median:0.407, p-value=0.063; Control baseline median:0.323, Control follow-up median:0.356, p-value=0.406). (G) Isotropy (Concussed baseline median:0.469, Concussed follow-up median:0.468, p-value=0.067; Control baseline median:0.371, Control follow-up median:0.416, p-value: 0.491).

Since our diffusion results indicated microstructural damage acutely after injury that appeared to recover in certain individuals by the time of clinical recovery, we determined how this affected global brain network structure in the SRC athletes. We used the GQI data to perform tractography and combined the resulting connectome with a population specific brain atlas to create a brain network representing the density of white matter tracts between brain regions for everyone at each time point (see Methods). We then calculated six measures of network structure as described in the Methods for each brain network to understand how global network organization might be impacted by injury and recovery.

Figure 4 presents the network measure comparisons at baseline. There were no significant differences between groups for the clustering coefficient, global efficiency, dissimilarity, spectral radius, synchronizability, or small-world propensity (Table S1). This is opposite to the diffusion measures, suggesting that any changes in brain network structure resulting from injury are not observable at the global scale at this timepoint and in this sample.

**Figure 4.**
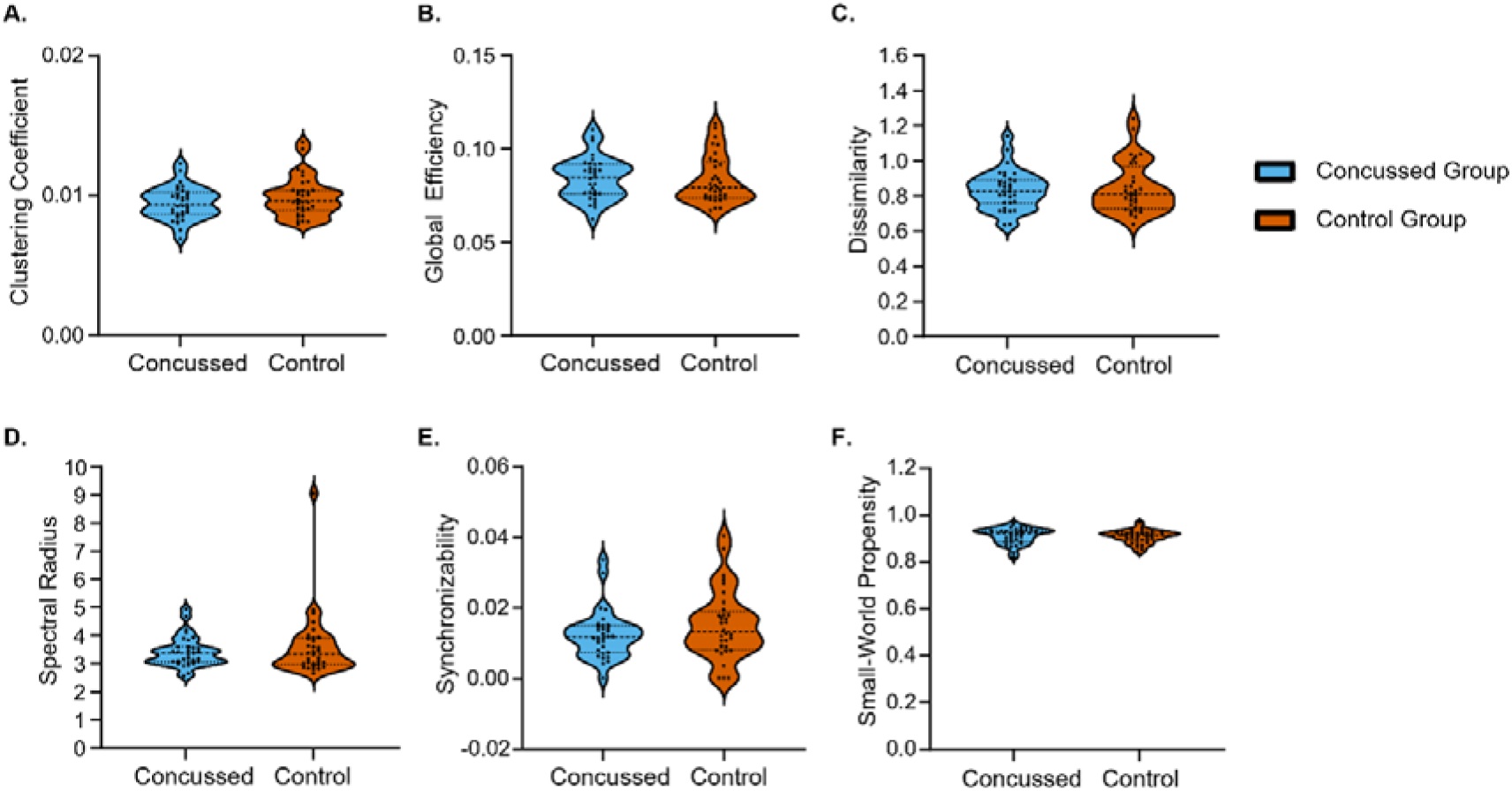
Network Measures Compared Between Groups at Baseline. Violin plots displaying the distributions of the network measures by group. (A) Clustering coefficient (Concussed median:0.009, Control median:0.010, p-value=0.328). (B) Global efficiency (Concussed median:0.085, Control median:0.079, p-value=0.490). (C) Dissimilarity (Concussed median:0.826, Control median:0.812, p-value=0.822). (D) Spectral radius (Concussed median:3.367, Control median:3.352, p-value=0.951). (E) Synchronizability (Concussed median:0.012, Control median:0.013, p-value=0.247). (F) Small-world propensity (Concussed median:0.920, Control median:0.914, p-value=0.525).

We also assessed network-level measures at the time of clinical recovery (Figure 5).

**Figure 5.**
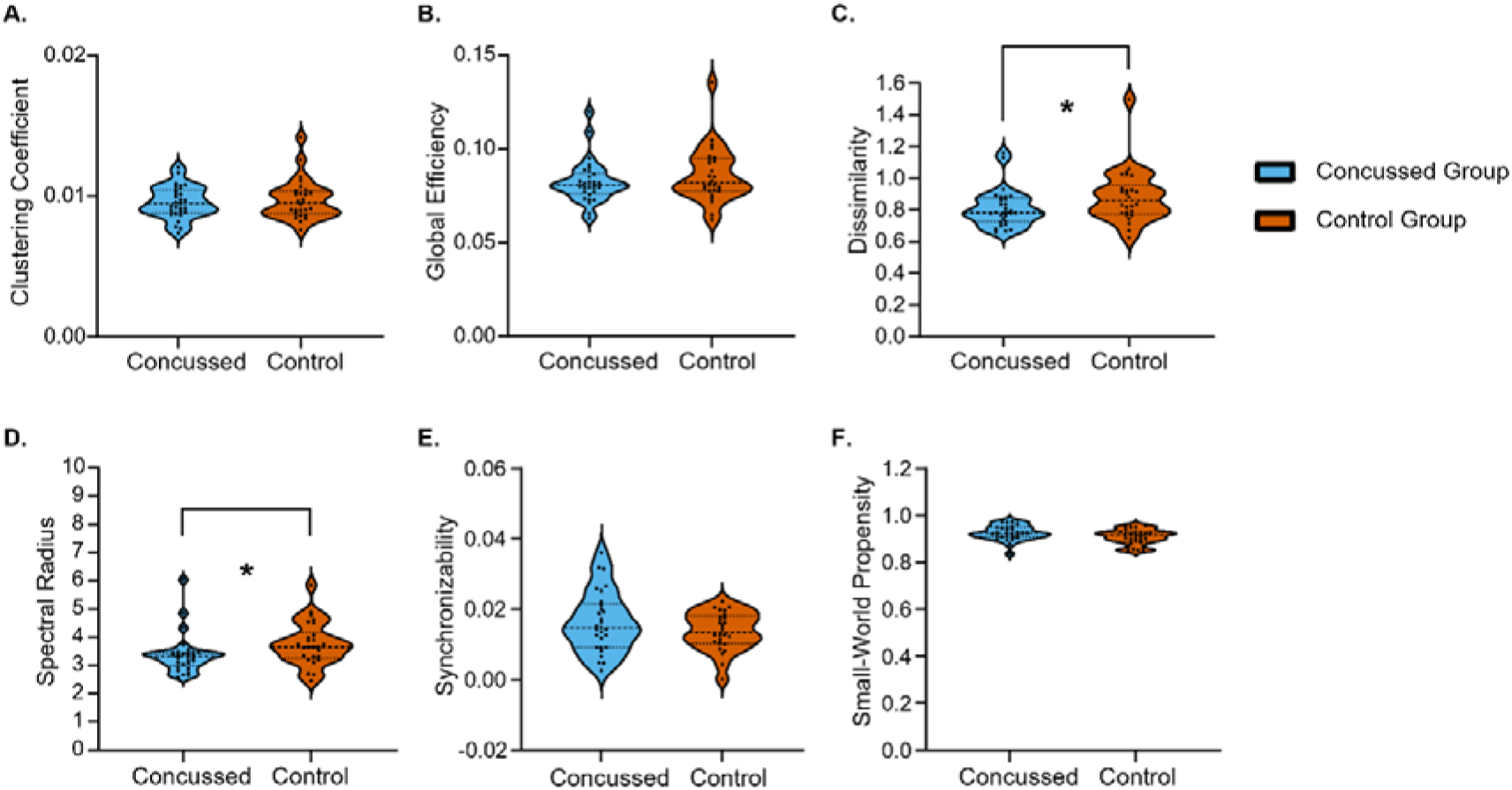
Network Measures Compared Between Groups at Follow-Up. Violin plots displaying the distributions of the network measures by group. (A) Clustering coefficient (Concussed median:0.009, Control median:0.010, p-value=0.953). (B) Global efficiency (Concussed median:0.081, Control median:0.082, p-value=0.361). (C) Dissimilarity (Concussed median:0.780, Control median:0.856, p-value=0.049). (D) Spectral radius (Concussed median:3.319, Control median:3.653, p-value=0.011). (E) Synchronizability (Concussed median:0.015, Control median:0.013, p-value=0.388). (F) Small-world propensity (Concussed median:0.924, Control median:0.915, p-value=0.104).

There were no significant differences observed between groups for clustering coefficient, global efficiency, synchronizability, or small-world propensity (Fig. 5A, B, E, F, Table S1). However, there was a significant difference between the concussed and control groups for dissimilarity and spectral radius (Fig. 5C and D, Table S1). This suggests that global changes in brain network structure are delayed in forming after the injury.

We also examined the change from baseline to follow-up for the network-level measures within the concussed and control groups (Figure 6). There were no significant differences observed between baseline and follow-up for the controls for clustering coefficient, global efficiency, dissimilarity, spectral radius, synchronizability, or small-world propensity (Table S2). The concussed group also showed no significant differences from baseline to follow-up for clustering coefficient, global efficiency, dissimilarity, or spectral radius (Fig. 6 A-D, Table S2). There were, however, significant increases from baseline to follow-up for synchronizability and small-world propensity (Fig. 6 E and F, Table S2). As with the previous analysis of diffusion measures, the data show a heterogeneous response, making it difficult to interpret these findings at the group level.

**Figure 6.**
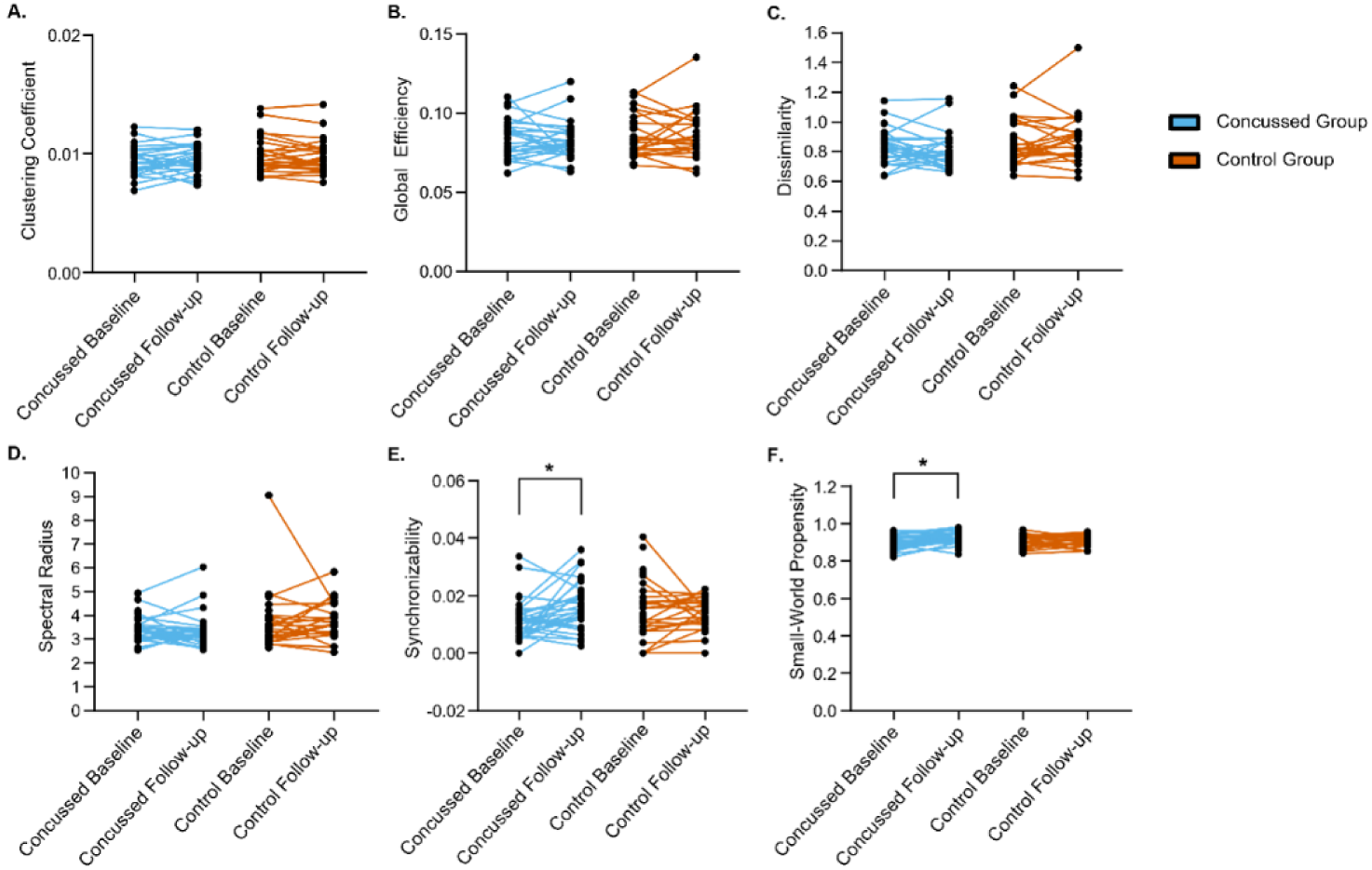
Network Measures Compared Between Groups for the Difference in Timepoints. Paired line plots displaying the change between baseline and follow-up for the network measures. (A) Clustering coefficient (Concussed baseline median:0.009, Concussed follow-up median:0.009, p-value=0.237; Control baseline median:0.010, Control follow-up median:0.010, p-value=0.264). (B) Global efficiency (Concussed baseline median:0.085, Concussed follow-up median:0.081, p-value=0.708 Control baseline median:0.079, Control follow-up median:0.082, p-value=0.684). (C) Dissimilarity (Concussed baseline median:0.826, Concussed follow-up median:0.780, p-value=0.437; Control baseline median:0.812, Control follow-up median:0.856, p-value=0.705). (D) Spectral radius (Concussed baseline median:3.367, Concussed follow-up median:3.319, p-value=0.635; Control baseline median:3.352, Control follow-up median:3.653, p-value=0.422). (E) Synchronizability (Concussed baseline median:0.012, Concussed follow-up median:0.015, p-value=0.025; Control baseline median:0.013, Control follow-up median:0.013, p-value=0.218). (F) Small-world propensity (Concussed baseline median:0.920, Concussed follow-up median:0.924, p-value=0.015; Control baseline median:0.914, Control follow-up median:0.915, p-value=0.565).

Taken together, these results indicate that microstructural damage occurs acutely after injury and may be resolved by clinical recovery, but network disruptions take longer to appear and persist beyond the point of clinical recovery. However, we also found that SRC are heterogenous, and these group-level analyses may mask subject-specific patterns of injury.

Given the observed heterogeneity, we performed an individual-level analysis using differential tractography, which calculates longitudinal white matter changes at the subject level (see the Methods) to identify specific track bundles that have changed between timepoints within a single individual.

Figure 7 presents differential tractography results for the concussed group, control group, and the concussed tracks with a frequency greater than the control-derived baseline. To better differentiate physiological change from changes that occurred due to injury, we calculated the frequency at which each identified track bundle occurred across individuals within each group. Tracks with a higher frequency in the concussed group than the controls were assumed to be associated with injury as opposed to chance variability. Results for QA and RDI are presented at the 25% threshold. The same analyses were performed for QA and RDI at the 30% and 35% thresholds (Tables S5, S6, S8, and S9). For QA at the 25% threshold there were 5/24 (20.8%) concussed and 2/24 (8.3%) control participants who had differential tractography results depicting changes in tracks over time. The concussed group aggregated tracks consisted of the commissure fiber in the corpus callosum, projection fibers at the brainstem and basal ganglia levels, and association fibers such as the longitudinal fasciculus, cingulum fibers, and occipital fasciculus (Fig. 7A). The control group aggregated tracks were largely similar with minor variability (Fig. 7B). The concussed tracks with a frequency above the controls included the corpus callosum forceps major and tapetum as well as the left corticospinal track at the brainstem level (Fig. 7C).

**Figure 7.**
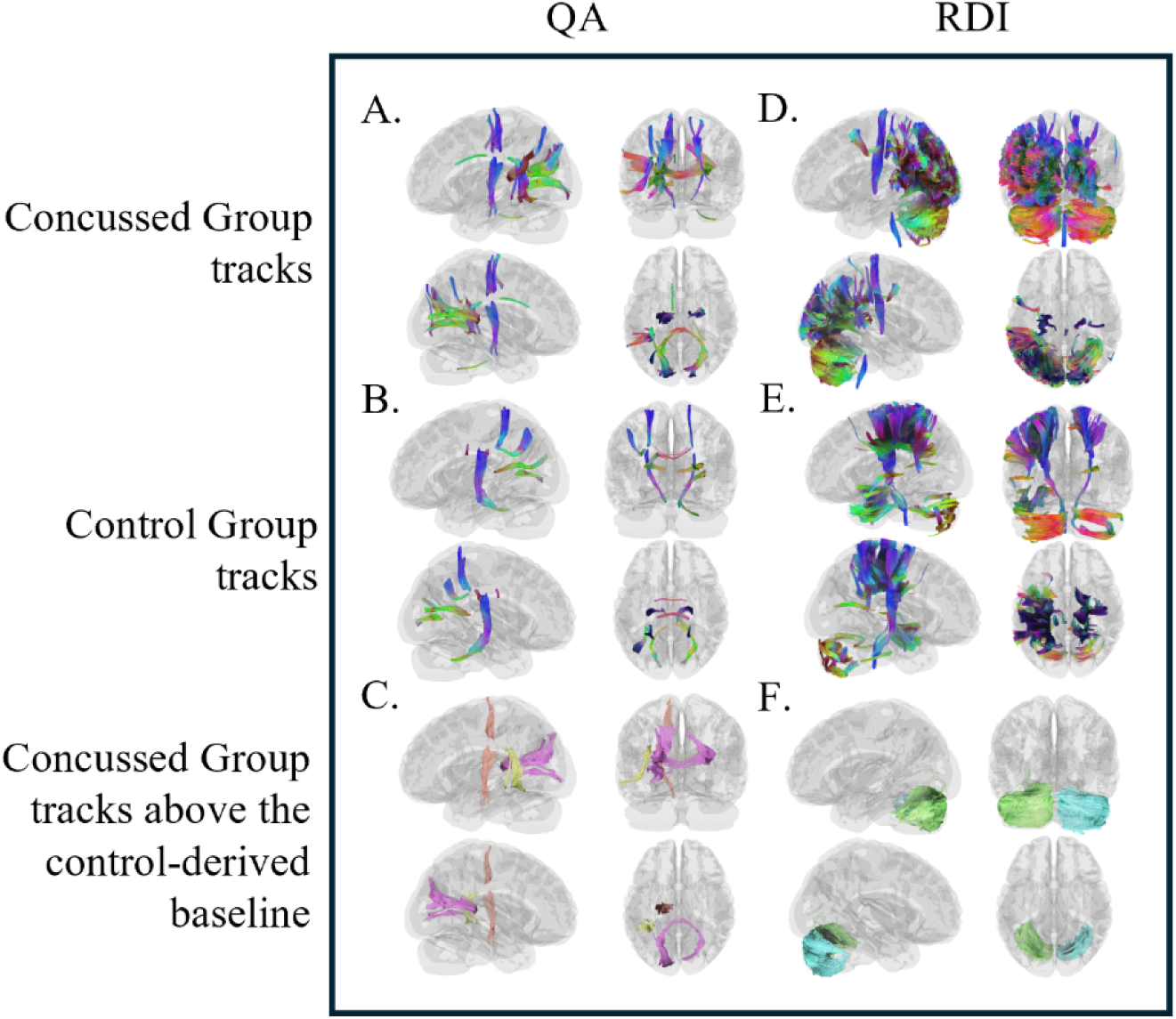
Differential tractography results aggregated by group for QA and RDI at the 25% threshold. (A) Differential tractography QA results for the concussed group (E.g., corpus callosum forceps major and tapetum, corticospinal tract, thalamic radiation, and corticotriatial tract). (B) QA results for the control group (E.g., corpus callosum forceps major and body, medial lemniscus, and corticopontine tract). (C) QA results for the concussed tracks with a frequency higher than the control-derived baseline (E.g., corpus callosum forceps major and tapetum and left corticospinal tract). (D) RDI results for the concussed group (E.g., cerebellum). (E) RDI results for the control group (E.g., cerebellum and corpus callosum). (F) RDI results for the concussed tracks with a frequency higher than the control-derived baseline (E.g., right and left cerebellum).

For RDI at the 25% threshold, there were 9/24 (37.5%) concussed and 8/24 (33.3%) control participants with differential tractography results. The concussed group aggregated tracks consisted of cerebellum fibers, association fibers including the longitudinal and occipital fasciculi, and brainstem projection fibers (Fig. 7D). Controls had many similar fibers present (Fig. 7E). The concussed tracks with a frequency above the controls included the left and right cerebellar tracks (Fig. 7F). The concussed group had patterns of damage localized to the cerebellum, corpus callosum, and corticospinal track.

We then examined the data stratified by sex. Figure 8 presents the differential tractography data stratified by sex for QA and RDI at the 25% thresholds. The 30% and 35% thresholds are presented in the supplemental materials (Tables S11, S12, S14, S15, S17, S18, S20, and S21). There were 4/14 (28.6%) concussed males and 1/15 (6.7%) control males with differential tractography results. The male concussed and control group for QA at 25% had similar tracks present including the corpus callosum, projection fibers at the brainstem and basal ganglia levels, and association fibers (Fig. 7A and B). All male concussed tracks had a frequency above the control-derived baseline (Fig. 7C). For females, only 1/10 (10.0%) concussed and 1/9 (11.1%) controls had differential tractography results. Female concussed athletes for QA at 25% had more variety in the fibers that showed change after differential tractography, which were similar to those seen in the male concussed group (Fig. 7D). The control females had a large majority of brainstem and basal ganglia projection fibers present (Fig. 7E), and the concussed females had no tracks with a greater frequency than the control baseline (Fig. 7F). This suggests that the changes in differential tractography after injury are almost exclusively occurring in male participants.

**Figure 8.**
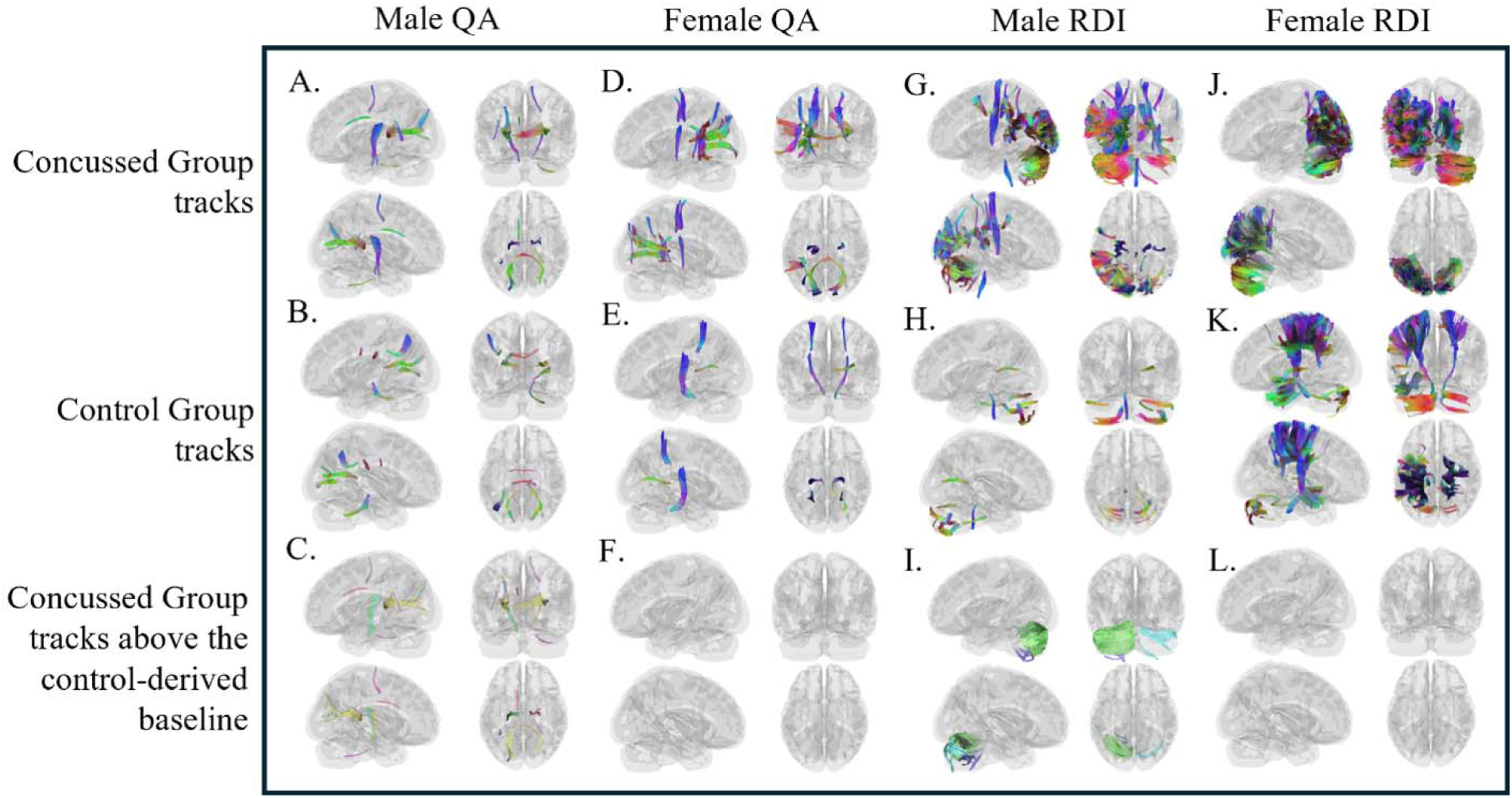
Differential tractography results stratified by sex for QA and RDI at the 25% threshold. (A) Male differential tractography QA results for the concussed group. (B) Male QA results for the control group. (C) Male QA results for the concussed tracks with a frequency higher than the control-derived baseline. (D) Female QA results for the concussed group. (E) Female QA results for the control group. (F) Female QA results for the concussed tracks with a frequency higher than the control-derived baseline. (G) Male differential tractography RDI results for the concussed group. (H) Male RDI results for the control group. (I) Male RDI results for the concussed tracks with a frequency higher than the control-derived baseline. (J) Female RDI results for the concussed group. (K) Female RDI results for the control group. (L) Female RDI results for the concussed tracks with a frequency higher than the control-derived baseline.

For RDI at 25% there were 6/14 (42.9%) concussed males and 4/15 (26.7%) control males with differential tractography results. The concussed males had tracks similar to control males, the majority which consisted of cerebellar fibers (Fig. 7G and H). The concussed males’ vermis and left and right cerebellar hemispheres had a frequency higher than control males (Fig 7I). The females had 3/10 (30.0%) concussed and 4/9 (44.4%) control participants with differential tractography results. The concussed and control females also had similar track profiles, but the concussed females had no tracks above the control females’ baseline frequency (Fig 7J-L). This confirmed that concussed males accounted for the patterns of damage seen in the differential tractography analysis stratified by group, whereas concussed females showed little white matter damage overall.

## 4. DISCUSSION

This study compared diffusion and network measures between concussed and control adolescent athletes and examined individual variability in brain structure stratified by group and sex.

### 4.1 Diffusion measures: Microstructural changes that are present acutely after injury resolve by clinical recovery

DTI measures have been studied extensively in concussion research.^13–15,47–49^ In this study, we found no differences in DTI measures between groups at baseline or follow-up. Some studies have corroborated this lack of DTI group difference after injury^50–52^, while others have reported differences after injury but not at recovery^53,54^, yet others have reported differences from injury that persisted beyond clinical recovery^55,56^. These discrepancies may be related to methodological differences among diffusion models.

This is the first study to our knowledge to investigate quantitively derived GQI measures longitudinally after adolescent SRC. GQI provides improved resolution of complex fiber architecture that may not be adequately captured by the single tensor DTI framework.

Specifically, we found that QA, RDI, and ISO were significantly increased in the concussed group at baseline, but not at follow-up. QA, RDI, and ISO are all quantitively derived GQI measures thought to reflect white matter structure (i.e., axonal density)^36,40^, inflammation (i.e., cellular density)^39,57^, and background, non-directional diffusion (i.e., cerebrospinal fluid or edema)^36^, respectively. The greater QA and RDI acutely after SRC may indicate changes in white matter structure and neuroinflammation. This is corroborated by an increase in non-directional or isotropic diffusion, ISO, compared to controls, which suggests increased extracellular water or possible edema due to SRC. At the follow-up timepoint, i.e. clinical recovery, the lack of differences suggests resolution of these microstructural changes associated with a return to pre-injury status as symptoms resolved.

### 4.2 Network measures: Network disruptions not present acutely after injury appeared by clinical recovery and likely persisted

At the network level, we found no differences at the baseline timepoint between the groups, but at follow-up the concussed group had significantly reduced dissimilarity and spectral radius compared to the control group. Dissimilarity provides information on how dissimilar structural connectivity matrices are for a group and spectral radius is indicative of activity spread in a network.^44,45^ These results suggest that at follow-up, the concussed groups’ brain networks were more similar to each other and had less capacity for information spread than the controls.

Consistent with our findings, prior work has observed a lack of network alterations acutely after injury, but emergent, widespread network alterations in mTBI at 1-year post-injury.^58,59^

We also observed within group differences for the network measures synchronizability and small-world propensity, which were increased in the concussed group at follow-up compared to baseline. Prior studies have also reported an increased small-worldness in pediatric, chronic mTBI populations^12^. This small-world topology, which reflects a balance of integration and segregation, is thought to be the innate structure of human brain networks.^60^ The increased small-world propensity at follow-up compared to baseline suggests that the concussed groups’ brain networks, which likely had been disrupted acutely, expressed changes at a slower rate such that they could be measured only at or after the resolution of clinical symptoms. Small-world topology is known to facilitate fast, stable brain dynamics, and synchronizability quantifies the ease with which a brain network can engages in coordinated, logical activity.^43,61^ Thus, greater small-world propensity and synchronizability in our concussed group suggests ongoing modified brain dynamics at the time of clinical recovery. Taken together, these results indicate that large-scale network reorganization may persist beyond normalization of microstructural changes and clinical symptom resolution. Like the difference from baseline to follow-up for the diffusion measures, the within group differences for the network measures were heterogeneous among participants. Future studies should confirm these results with larger sample sizes.

### 4.3 Differential tractography: Concussed group shows consistent damage, primarily in males

One obstacle to clinical dMRI adoption is that current literature reports group level results; the findings cannot be translated reliably to the subject level.^63^ In this study, however, we found a consistent pattern of injured white matter tracts (reduced QA and RDI) in the individual subjects in the concussed group, and when stratified by sex, we found it only in males; females had no damaged white matter tracks when compared to the control-derived noise baseline.

Within the patterns of damaged white matter tracks in the concussed group, we consistently found that the corpus callosum was present. The corpus callosum, a large tract connecting the right and left hemispheres, is important for cognitive function and has been shown to be injured following concussion.^64,65^ We also found QA and RDI changes in the corticospinal tract at the brainstem level in our concussed group. The corticospinal tracts are critical for motor control, but damage here may also result in headaches, balance problems, or cognitive deficits.^66,67^ The brainstem is known to be susceptible to shearing forces and is thought to be more vulnerable than other brain regions to the effects of SRC.^22,24,68,69^

In addition to the brain regions known to be damaged after a concussion, we also found notable sex differences. One diffusion imaging study of concussed female athletes 6-months from injury and controls found MD and RD differences in the corpus callosum but nothing in the corticospinal tract between groups.^64^ Clinical research has shown that female athletes tend to have greater symptom burden (number and severity) and sometimes longer recovery times versus males, whereas neuroimaging research has been more mixed^26^ Our concussed females athletes reported a similar total symptom score of 36.11 (max = 126) compared to males (35.50), but did not show SRC white matter alterations. Taken together our results suggest the possibility of sex-specific patterns of post-injury reorganization or susceptibility to SRC.

QA and RDI increases were tested in addition to decreases; however, these increases were not consistently detected across thresholds. This limits the interpretability of the increases in QA and RDI within the current tractography framework employed here. This lack of stable increase findings may suggest that the detected tractography changes were primarily driven by decreases in QA and RDI rather than bidirectional alterations. Future studies should further explore the possibility of increases in a larger sample size.

### 4.4 Study Limitations

This study presents novel findings of structural brain network changes at the individual level post-concussion, but it is not without limitations. Our sample consists of adolescent athletes with typical, non-delayed recovery and may not be generalizable to other populations, other mechanisms of injury, or to those who have symptoms that persist for months or years. We did not account for developmental changes between our sample participants in this study. We also have no objective information or detailed self-reporting of how the injury occurred to hypothesize regions of the brain that may be damaged after SRC; thus we assessed whole brain differences. Our concussed participants reported more prior concussions when compared to the controls. This is a well-known finding in the SRC literature, where having had a concussion increases the chance of being diagnosed with another one.^70^ However, we did not control our analyses for these demographic differences. When assessing sex differences, we did not collect subjective or objective hormonal or menstrual cycle data from female participants. Future studies should aim to collect injury details, whether self-reported or objective (e.g., video footage of the game or accelerometer helmet data), hormonal data from female participants, and a larger sample size that can be divided into developmental subgroups.

Despite these limitations, the early group-level white matter microstructure damage and the delayed, and potentially persisting, network reorganization we observed remain important findings. Future research should corroborate these findings with clinical symptoms and outcomes in males and females separately in a larger sample, and account for clinical symptoms, time to recovery, and relation to hormonal status. Our results indicate that more individualized approaches should be developed and analyzed to fully understand the heterogenous nature of SRC.

## Data and Code Availability

The data and code used in this study are available from the corresponding author upon reasonable request.

## Author Contributions

E.V.C.: Formal Analysis, Investigation, Project Administration, Visualization, Writing – original draft; M.N.H.: Investigation, Project Administration, Writing – review and editing; F.S.: Investigation, Project Administration, Writing – review and editing; J.J.L.: Conceptualization, Funding Acquisition, Project Administration, Supervision, Writing – review and editing; J.C.M.: Investigation, Writing – review and editing; S.M.: Conceptualization, Resources, Project Administration, Supervision, Writing – review and editing.

