## Supplemental file for "Sport-Related Concussion in Adolescent Athletes is Associated with Acute White Matter Alterations, Delayed Network Changes, and Individual-Level Injury Patterns"

**Table S1. Diffusion/GQI and Network Measure Results Summary for Baseline and Follow-Up**

| **Model** | **Response Variable** | **Concussed Group Mean** | **Control Group Mean** | **W** | **Effect Size (95% Confidence Interval)** | ***p*-value** |
|---|---|---|---|---|---|---|
| Diffusion/GQI Baseline | | | | | | |
| | Fractional Anisotropy | 0.4309574 | 0.4329172 | 576 | 0.139 (0.007, 0.380) | 0.2765 |
| | Mean Diffusivity | 0.7918925 | 0.8037202 | 438 | 0.100 (0.004, 0.330) | 0.4319 |
| | Radial Diffusivity | 0.5951847 | 0.6007271 | 414 | 0.142 (0.009, 0.390) | 0.2645 |
| | Axial Diffusivity | 1.207511 | 1.201266 | 478 | 0.031 (0.005, 0.300) | 0.8112 |
| | Quantitative Anisotropy | 0.2599666 | 0.2063979 | 680 | 0.319 (0.100, 0.550) | **0.0120** |
| | Restricted Diffusion Imaging | 0.4105486 | 0.3233440 | 658 | 0.281 (0.030, 0.520) | **0.0257** |
| | Isotropy | 0.4692100 | 0.3707407 | 650 | 0.267 (0.030, 0.490) | **0.0342** |
| Diffusion/GQI Follow-up | | | | | | |
| | Fractional Anisotropy | 0.4339790 | 0.4308476 | 436 | 0.134 (0.009, 0.400) | 0.3268 |
| | Mean Diffusivity | 0.8017202 | 0.8048531 | 323 | 0.123 (0.007, 0.370) | 0.3700 |
| | Radial Diffusivity | 0.6013405 | 0.6033070 | 317 | 0.136 (0.009, 0.390) | 0.3185 |
| | Axial Diffusivity | 1.197070 | 1.205978 | 321 | 0.127 (0.005, 0.380) | 0.3523 |
| | Quantitative Anisotropy | 0.2586494 | 0.2203904 | 396 | 0.043 (0.005, 0.300) | 0.757 |
| | Restricted Diffusion Imaging | 0.4069236 | 0.3560568 | 380 | 0.007 (0.005, 0.300) | 0.9667 |
| | Isotropy | 0.4683351 | 0.4162677 | 374 | 0.007 (0.005, 0.300) | 0.9667 |
| Network Baseline | | | | | | |
| | Clustering Coefficient | 0.009342130 | 0.009621097 | 424 | 0.125 (0.005, 0.350) | 0.328 |
| | Global Efficiency | 0.08467206 | 0.07910459 | 547 | 0.088 (0.004, 0.340) | 0.4902 |
| | Dissimilarity | 0.8255982 | 0.8124128 | 513 | 0.029 (0.005, 0.290) | 0.8218 |

| | | | | | | |
|---|---|---|---|---|---|---|
| | Spectral Radius | 3.367224 | 3.351839 | 501 | 0.009 (0.004, 0.290) | 0.951 |
| | Synchronizability | 0.01190780 | 0.01326942 | 411 | 0.147 (0.005, 0.380) | 0.2472 |
| | Small-World Propensity | 0.9203898 | 0.9138679 | 543 | 0.081 (0.005, 0.340) | 0.5252 |
| Network Follow-up | | | | | | |
| | Clustering Coefficient | 0.009487290 | 0.009504938 | 373 | 0.009 (0.002, 0.310) | 0.9533 |
| | Global Efficiency | 0.0807153 | 0.0817797 | 322 | 0.125 (0.005, 0.380) | 0.3611 |
| | Dissimilarity | 0.7802914 | 0.8564457 | 260 | 0.266 (0.020, 0.490) | **0.0490** |
| | Spectral Radius | 3.319185 | 3.653268 | 227 | 0.341 (0.090, 0.570) | **0.0109** |
| | Synchronizability | 0.01484230 | 0.01343482 | 429 | 0.118 (0.007, 0.370) | 0.3883 |
| | Small-World Propensity | 0.9240417 | 0.9150668 | 474 | 0.220 (0.020, 0.460) | 0.1041 |

**Table S2. Diffusion/GQI and Network Measure Statistical Results for the Difference in Timepoints**

| Group and Model | Response Variable | Baseline Median | Follow-Up Median | V | Effect Size (95% Confidence Interval) | *p*-value |
|---|---|---|---|---|---|---|
| Concussed Diffusion/GQI Difference | | | | | | |
| | Fractional Anisotropy | 0.4309574 | 0.433979 | 238 | 0.360 (-0.070, 0.670) | 0.1165 |
| | Mean Diffusivity | 0.7918925 | 0.8017202 | 134 | -0.240 (-0.590, 0.200) | 0.3032 |
| | Radial Diffusivity | 0.5951847 | 0.6013405 | 119 | -0.320 (-0.650, 0.110) | 0.1574 |
| | Axial Diffusivity | 1.207511 | 1.19707 | 164 | -0.070 (-0.470, 0.360) | 0.7835 |
| | Quantitative Anisotropy | 0.2599666 | 0.2586494 | 251 | 0.430 (0.020, 0.720) | 0.05585 |
| | Restricted Diffusion Imaging | 0.4105486 | 0.4069236 | 249 | 0.420 (0.010, 0.710) | 0.06303 |
| | Isotropy | 0.46921 | 0.4683351 | 248 | 0.410 (0.000, 0.710) | 0.06688 |
| Control Diffusion/GQI Difference | | | | | | |
| | Fractional Anisotropy | 0.4329172 | 0.4308476 | 170 | 0.130 (-0.310, 0.530) | 0.5838 |
| | Mean Diffusivity | 0.8037202 | 0.8048531 | 143 | -0.050 (-0.470, 0.390) | 0.8553 |
| | Radial Diffusivity | 0.6007271 | 0.603307 | 140 | -0.070 (-0.480, 0.370) | 0.7898 |
| | Axial Diffusivity | 1.201266 | 1.205978 | 150 | 0.000 (-0.430, 0.430) | 0.9999 |
| | Quantitative Anisotropy | 0.2063979 | 0.2203904 | 113 | -0.250 (-0.610, 0.200) | 0.3029 |
| | Restricted Diffusion Imaging | 0.323344 | 0.3560568 | 120 | -0.200 (-0.580, 0.250) | 0.4061 |
| | Isotropy | 0.3707407 | 0.4162677 | 125 | -0.170 (-0.550, 0.280) | 0.4908 |
| Concussed Network Difference | | | | | | |
| | Clustering Coefficient | 0.00934213 | 0.00948729 | 128 | -0.270 (-0.620, 0.160) | 0.237 |
| | Global Efficiency | 0.08467206 | 0.0807153 | 191 | 0.090 (-0.340, 0.480) | 0.7078 |

| | | | | | | |
|---|---|---|---|---|---|---|
| | Dissimilarity | 0.8255982 | 0.7802914 | 207 | 0.180 (-0.250, 0.550) | 0.4374 |
| | Spectral Radius | 3.367224 | 3.319185 | 195 | 0.110 (-0.320, 0.500) | 0.6348 |
| | Synchronizability | 0.0119078 | 0.0148423 | 88 | -0.500 (-0.760, -0.110) | **0.02535** |
| | Small-World Propensity | 0.9203898 | 0.9240417 | 81 | -0.540 (-0.780, -0.160) | **0.01513** |
| Control Network Difference | | | | | | |
| | Clustering Coefficient | 0.009621097 | 0.009504938 | 190 | 0.270 (-0.180, 0.620) | 0.2643 |
| | Global Efficiency | 0.07910459 | 0.0817797 | 135 | -0.100 (-0.510, 0.340) | 0.684 |
| | Dissimilarity | 0.8124128 | 0.8564457 | 136 | -0.090 (-0.500, 0.350) | 0.7048 |
| | Spectral Radius | 3.351839 | 3.653268 | 121 | -0.190 (-0.570, 0.260) | 0.4223 |
| | Synchronizability | 0.01326942 | 0.01343482 | 106 | -0.290 (-0.640, 0.150) | 0.2182 |
| | Small-World Propensity | 0.9138679 | 0.9150668 | 129 | -0.140 (-0.540, -0.310) | 0.5646 |

**Table S3. Full List of Concussed and Control Tracts from Differential Tractography for QA 0.25**

| Concussed | Control |
|---|---|
| Commissure: Corpus Callosum Forceps Major | Commissure: Corpus Callosum Forceps Major |
| Brainstem: Left Corticospinal Tract | Commissure: Corpus Callosum Body |
| Commissure: Corpus Callosum Tapetum | Brainstem: Right Medial Lemniscus |
| Brainstem: Right Corticospinal Tract | Brainstem: Right Occipital Corticopontine Tract |
| Brainstem: Left Parietal Corticopontine Tract | Association: Right Inferior Fronto-Occipital Fasciculus |
| Basal Ganglia: Right Posterior Thalamic Radiation | Association: Left Superior Longitudinal Fasciculus |
| Basal Ganglia: Left Posterior Thalamic Radiation | Cerebellum: Middle Cerebellar Peduncle |
| Basal Ganglia: Right Superior Corticostriatal Tract | Brainstem: Right Non-Decussating Dentatorubrothalamic Tract |
| Basal Ganglia: Right Posterior Corticostriatal Tract | Association: Right Inferior Longitudinal Fasciculus |
| Commissure: Corpus Callosum Body | Commissure: Occipital Anterior Commissure |
| Association: Right Inferior Longitudinal Fasciculus | Brainstem: Left Occipital Corticopontine Tract |
| Association: Left Inferior Longitudinal Fasciculus | Basal Ganglia: Right Posterior Corticostriatal Tract |
| Association: Right Inferior Fronto-Occipital Fasciculus | Basal Ganglia: Right Optic Radiation |
| Association: Left Inferior Fronto-Occipital Fasciculus | Cerebellum: Superior Cerebellar Peduncle |
| Association: Left Arcuate Fasciculus | Brainstem: Left Corticospinal Tract |
| Brainstem: Right Non-Decussating Dentatorubrothalamic Tract | Brainstem: Right Corticospinal Tract |
| Brainstem: Left Non-Decussating Dentatorubrothalamic Tract | Brainstem: Right Parietal Corticopontine Tract |
| Brainstem: Right Medial Lemniscus | Brainstem: Left Parietal Corticopontine Tract |
| Brainstem: Left Medial Lemniscus | Brainstem: Left Medial Lemniscus |
| Brainstem: Right Parietal Corticopontine Tract | Basal Ganglia: Left Superior Thalamic Radiation |
| Brainstem: Left Occipital Corticopontine Tract | Association: Right Extreme Capsule |
| Basal Ganglia: Left Superior Thalamic Radiation | Basal Ganglia: Left Posterior Corticostriatal Tract |
| Basal Ganglia: Right Optic Radiation | Basal Ganglia: Right Superior Thalamic Radiation |
| Basal Ganglia: Left Optic Radiation | Association: Left Extreme Capsule |
| Basal Ganglia: Left Superior Corticostriatal Tract | Basal Ganglia: Left Superior Corticostriatal Tract |
| Basal Ganglia: Left Posterior Corticostriatal Tract | Association: Left Middle Longitudinal Fasciculus |

| |
|---|
| Commissure: Occipital Anterior Commissure<br>Cerebellum: Middle Cerebellar Peduncle<br>Association: Left Parietal Aslant Tract<br>Association: Left Middle Longitudinal Fasciculus<br>Association: Left Frontal Parietal Cingulum<br>Association: Left Frontal Parahippocampal Cingulum |

*Concussed tracts with a frequency above the control tracts are highlighted in red.*

**Table S4. Full List of Concussed and Control Tracts from Differential Tractography for QA 0.30**

| Concussed | Control |
|---|---|
| Commissure: Corpus Callosum Forceps Major | Commissure: Corpus Callosum Forceps Major |
| Basal Ganglia: Left Posterior Thalamic Radiation | Brainstem: Left Parietal Corticopontine Tract |
| Brainstem: Left Corticospinal Tract | Brainstem: Left Corticospinal Tract |
| Commissure: Corpus Callosum Tapetum | Brainstem: Left Medial Lemniscus |
| Association: Left Arcuate Fasciculus | Commissure: Corpus Callosum Body |
| Association: Right Inferior Longitudinal Fasciculus | Association: Left Superior Longitudinal Fasciculus |
| Association: Left Parietal Aslant Tract | Basal Ganglia: Right Posterior Corticostriatal Tract |
| Basal Ganglia: Right Posterior Corticostriatal Tract | |
| Commissure: Occipital Anterior Commissure | |
| Basal Ganglia: Left Optic Radiation | |
| Brainstem: Left Occipital Corticopontine Tract | |
| Association: Right Inferior Fronto-Occipital Fasciculus | |
| Basal Ganglia: Corticostriatal Tract | |
| Association: Left Inferior Longitudinal Fasciculus | |

*Concussed tracts with a frequency above the control tracts are highlighted in red. 4/24 (16.7%) of concussed participants and 2/24 (8.3%) of control participants had differential tractography results.*

**Table S5. Full List of Concussed and Control Tracts from Differential Tractography for QA 0.35**

| **Concussed** | **Control** |
|---|---|
| Commissure: Corpus Callosum Forceps Major<br>Association: Right Inferior Longitudinal Fasciculus<br>Commissure: Corpus Callosum Tapetum<br>Basal Ganglia: Left Posterior Thalamic Radiation<br>Basal Ganglia: Right Posterior Corticostriatal Tract | N/A |

*Concussed tracts with a frequency above the control tracts are highlighted in red. N/A indicates that no participants in that group had results. 1/24 (4.2%) of concussed participants and 0/24 (0.0%) of control participants had differential tractography results.*

**Table S6. Full List of Concussed and Control Tracts from Differential Tractography for RDI 0.25**

| Concussed | Control |
|---|---|
| Cerebellum: Right Cerebellum | Cerebellum: Left Cerebellum |
| Cerebellum: Left Cerebellum | Cerebellum: Right Cerebellum |
| Cerebellum: Vermis | Cerebellum: Vermis |
| Commissure: Corpus Callosum Forceps Major | Commissure: Corpus Callosum Forceps Major |
| Association: Left Inferior Fronto-Occipital Fasciculus | Brainstem: Left Medial Lemniscus |
| Commissure: Corpus Callosum Body | Cerebellum: Middle Cerebellar Peduncle |
| Association: Left Inferior Longitudinal Fasciculus | Brainstem: Left Corticospinal Tract |
| Association: Right Inferior Longitudinal Fasciculus | Brainstem: Left Medial Forebrain Bundle |
| Association: Right Inferior Fronto-Occipital Fasciculus | Association: Left Superior Longitudinal Fasciculus |
| Commissure: Occipital Anterior Commissure | Brainstem: Right Corticospinal Tract |
| Cerebellum: Middle Cerebellar Peduncle | Basal Ganglia: Left Superior Corticostriatal Tract |
| Association: Right Vertical Occipital Fasciculus | Commissure: Corpus Callosum Body |
| Association: Left Parietal Aslant Tract | Association: Left Extreme Capsule |
| Association: Left Vertical Occipital Fasciculus | Association: Left Inferior Longitudinal Fasciculus |
| Association: Left Arcuate Fasciculus | Brainstem: Left Parietal Corticopontine Tract |
| Brainstem: Left Parietal Corticopontine Tract | Association: Left Uncinate Fasciculus |
| Basal Ganglia: Left Superior Thalamic Radiation | Basal Ganglia: Right Superior Corticostriatal Tract |
| Basal Ganglia: Left Optic Radiation | Association: Left Parietal Aslant Tract |
| Basal Ganglia: Left Posterior Thalamic Radiation | Association: Left Arcuate Fasciculus |
| Association: Left Superior Longitudinal Fasciculus | Brainstem: Right Medial Lemniscus |
| Basal Ganglia: Left Posterior Corticostriatal Tract | Basal Ganglia: Left Superior Thalamic Radiation |
| Basal Ganglia: Right Posterior Corticostriatal Tract | Brainstem: Right Parietal Corticopontine Tract |
| Commissure: Corpus Callosum Tapetum | Commissure: Temporal Anterior Commissure |
| Brainstem: Left Occipital Corticopontine Tract | Association: Left Middle Longitudinal Fasciculus |
| Basal Ganglia: Right Optic Radiation | Basal Ganglia: Right Superior Thalamic Radiation |
| Association: Left Superior Longitudinal Fasciculus | Brainstem: Right Non-Decussating Dentatorubrothalamic Tract |
| Basal Ganglia: Right Posterior Thalamic Radiation | Association: Left Hippocampus Alveus |
| | Association: Right Extreme Capsule |
| | Brainstem: Non-Decussating Dentatorubrothalamic Tract |
| | Basal Ganglia: Left Posterior Corticostriatal Tract |

| | |
|---|---|
| Association: Left Middle Longitudinal Fasciculus<br>Brainstem: Right Occipital Corticopontine Tract<br>Association: Right Parietal Aslant Tract<br>Brainstem: Right Reticular Tract<br>Brainstem: Right Medial Forebrain Bundle<br>Brainstem: Left Reticular Tract<br>Cerebellum: Right Inferior Cerebellar Peduncle<br>Brainstem: Left Corticospinal Tract<br>Brainstem: Right Corticospinal Tract<br>Basal Ganglia: Left Superior Corticostriatal Tract<br>Brainstem: Right Medial Lemniscus<br>Brainstem: Left Medial Lemniscus<br>Brainstem: Right Parietal Corticopontine Tract<br>Association: Left Extreme Capsule<br>Association: Left Frontal Aslant Tract<br>Association: Right Superior Longitudinal Fasciculus Cingulum<br>Basal Ganglia: Right Superior Corticostriatal Tract<br>Basal Ganglia: Right Superior Thalamic Radiation<br>Brainstem: Left Non-Decussating Dentatorubrothalamic Tract | Association: Left Superior Longitudinal Fasciculus<br>Association: Left Frontal Aslant Tract<br>Brainstem: Left Occipital Corticopontine Tract<br>Brainstem: Right Frontal Corticopontine Tract<br>Basal Ganglia: Left Posterior Thalamic Radiation<br>Association: Right Superior Longitudinal Fasciculus Cingulum<br>Brainstem: Right Medial Forebrain Bundle<br>Association: Right Frontal Aslant Tract<br>Association: Right Superior Longitudinal Fasciculus<br>Commissure: Corpus Callosum Tapetum<br>Association: Left Superior Longitudinal Fasciculus Cingulum<br>Brainstem: Right Occipital Corticopontine Tract<br>Commissure: Occipital AnteriorCommissure<br>Association: Left Inferior Fronto-Occipital Fasciculus<br>Brainstem: Corticobulbar Tract |

*Concussed tracts with a frequency above the control tracts are highlighted in red.*

**Table S7. Full List of Concussed and Control Tracts from Differential Tractography for RDI 0.30**

| Concussed | Control |
|---|---|
| Cerebellum: Left Cerebellum | Cerebellum: Left Cerebellum |
| Cerebellum: Right Cerebellum | Cerebellum: Vermis |
| Cerebellum: Vermis | Cerebellum: Right Cerebellum |
| Commissure: Corpus Callosum Forceps Major | Cerebellum: Middle Cerebellar Peduncle |
| Association: Left Inferior Fronto-Occipital Fasciculus | Association: Left Superior Longitudinal Fasciculus |
| Association: Left Inferior Longitudinal Fasciculus | Brainstem: Left Corticospinal Tract |
| Association: Right Inferior Longitudinal Fasciculus | Association: Left Extreme Capsule |
| Basal Ganglia: Left Posterior Thalamic Radiation | Brainstem: Left Parietal Corticopontine Tract |
| Association: Left Vertical Occipital Fasciculus | Commissure: Corpus Callosum Body |
| Basal Ganglia: Right Posterior Corticostriatal Tract | Brainstem: Left Medial Lemniscus |
| Brainstem: Left Occipital Corticopontine Tract | Association: Left Middle Longitudinal Fasciculus |
| Cerebellum: Middle Cerebellar Peduncle | Basal Ganglia: Left Superior Thalamic Radiation |
| Basal Ganglia: Right Posterior Thalamic Radiation | Brainstem: Right Corticospinal Tract |
| Association: Left Parietal Aslant Tract | Basal Ganglia: Left Superior Corticostriatal Tract |
| Basal Ganglia: Left Posterior Corticostriatal Tract | Brainstem: Right Parietal Corticopontine Tract |
| Association: Right Inferior Fronto-Occipital Fasciculus | Brainstem: Right Medial Lemniscus |
| Basal Ganglia: Left Optic Radiation | Basal Ganglia: Left Posterior Thalamic Radiation |
| Brainstem: Left Parietal Corticopontine Tract | Basal Ganglia: Left Posterior Corticostriatal Tract |
| Commissure: Occipital Anterior Commissure | Association: Right Extreme Capsule |
| Association: Left Superior Longitudinal Fasciculus | Basal Ganglia: Right Superior Corticostriatal Tract |
| Commissure: Corpus Callosum Body | Brainstem: Left Non-Decussating Dentatorubrothalamic Tract |
| Association: Left Middle Longitudinal Fasciculus | Brainstem: Right Non-Decussating Dentatorubrothalamic Tract |
| Association: Right Vertical Occipital Fasciculus | |
| Basal Ganglia: Right Optic Radiation | |
| Association: Left Arcuate Fasciculus | |
| Cerebellum: Right Inferior Cerebellar Peduncle | |

*Concussed tracts with a frequency above the control tracts are highlighted in red. 9/24 (37.5%) of concussed participants and 5/24 (20.8%) of control participants had differential tractography results.*

**Table S8. Full List of Concussed and Control Tracts from Differential Tractography for RDI 0.35**

| Concussed | Control |
|---|---|
| Cerebellum: Left Cerebellum<br>Cerebellum: Right Cerebellum<br>Cerebellum: Middle Cerebellar Peduncle<br>Association: Left Vertical Occipital Fasciculus<br>Association: Inferior Fronto-Occipital Fasciculus<br>Commissure: Corpus Callosum Forceps Major<br>Association: Left Inferior Longitudinal Fasciculus<br>Basal Ganglia: Left Posterior Corticostriatal Tract<br>Basal Ganglia: Left Optic Radiation<br>Basal Ganglia: Left Posterior Thalamic Radiation<br>Brainstem: Left Occipital Corticopontine Tract<br>Commissure: Occipital Anterior Commissure | Cerebellum: Left Cerebellum<br>Cerebellum: Right Cerebellum<br>Cerebellum: Middle Cerebellar Peduncle<br>Cerebellum: Vermis |

*Concussed tracts with a frequency above the control tracts are highlighted in red. 6/24 (25.0%) of concussed participants and 4/24 (16.7%) of control participants had differential tractography results.*

**Table S9. Full List of Male Concussed and Male Control Tracts from Differential Tractography for QA 0.25**

| Concussed | Control |
|---|---|
| Commissure: Corpus Callosum Forceps Major | Commissure: Corpus Callosum Forceps Major |
| Brainstem: Left Corticospinal Tract | Commissure: Corpus Callosum Body |
| Commissure: Corpus Callosum Tapetum | Association: Right Inferior Fronto-Occipital Fasciculus |
| Brainstem: Left Parietal Corticopontine Tract | Association: Left Superior Longitudinal Fasciculus |
| Basal Ganglia: Right Superior Corticostriatal Tract | Cerebellum: Middle Cerebellar Peduncle |
| Brainstem: Right Corticospinal Tract | Brainstem: Right Non-Decussating Dentatorubrothalamic Tract |
| Basal Ganglia: Right Optic Radiation | Association: Right Inferior Longitudinal Fasciculus |
| Association: Left Frontal Parietal Cingulum | Brainstem: Right Medial Lemniscus |
| Commissure: Corpus Callosum Body | Commissure: Occipital Anterior Commissure |
| Basal Ganglia: Right Posterior Corticostriatal Tract | Brainstem: Left Occipital Corticopontine Tract |
| Association: Left Frontal Parahippocampal Cingulum | Basal Ganglia: Right Posterior Corticostriatal Tract |
| Association: Right Inferior Longitudinal Fasciculus | Brainstem: Right Occipital Corticopontine Tract |
| Association: Right Inferior Fronto-Occipital Fasciculus | Basal Ganglia: Right Optic Radiation |
| Cerebellum: Middle Cerebellar Peduncle | Cerebellum: Superior Cerebellar Peduncle |
| Basal Ganglia: Left Posterior Thalamic Radiation | |
| Association: Left Inferior Longitudinal Fasciculus | |
| Association: Left Inferior Fronto-Occipital Fasciculus | |
| Association: Left Arcuate Fasciculus | |

*Concussed tracts with a frequency above the control tracts are highlighted in red.*

**Table S10. Full List of Male Concussed and Male Control Tracts from Differential Tractography for QA 0.30**

| Concussed | Control |
|---|---|
| Commissure: Corpus Callosum Forceps Major | Commissure: Corpus Callosum Forceps Major |
| Brainstem: Left Corticospinal Tract | Commissure: Corpus Callosum Body |
| Basal Ganglia: Left Posterior Thalamic Radiation | Association: Left Superior Longitudinal Fasciculus |
| Association: Left Inferior Longitudinal Fasciculus | Basal Ganglia: Right Posterior Corticostriatal Tract |

*Concussed tracts with a frequency above the control tracts are highlighted in red. 3/14 (21.4%) of male concussed participants and 1/15 (6.7%) of male control participants had differential tractography results.*

**Table S11. Full List of Male Concussed and Male Control Tracts from Differential Tractography for QA 0.35**

| Concussed | Control |
|---|---|
| N/A | N/A |

*Concussed tracts with a frequency above the control tracts are highlighted in red. N/A indicates that no participants in that group had results. 0/14 (0.0%) of male concussed participants and 1/15 (6.7%) of male control participants had differential tractography results.*

**Table S12. Full List of Female Concussed and Female Control Tracts from Differential Tractography for QA 0.25**

| Concussed | Control |
|---|---|
| Commissure: Corpus Callosum Forceps Major<br>Association: Left Arcuate Fasciculus<br>Commissure: Corpus Callosum Tapetum<br>Association: Left Parietal Aslant Tract<br>Brainstem: Left Corticospinal Tract<br>Basal Ganglia: Left Superior Corticostriatal Tract<br>Association: Right Inferior Longitudinal Fasciculus<br>Commissure: Occipital Anterior Commissure<br>Basal Ganglia: Left Optic Radiation<br>Association: Left Middle Longitudinal Fasciculus<br>Commissure: Corpus Callosum Body<br>Basal Ganglia: Right Posterior Corticostriatal Tract<br>Association: Right Inferior Fronto-Occipital Fasciculus<br>Basal Ganglia: Left Posterior Corticostriatal Tract<br>Brainstem: Right Non-Decussating Dentatorubrothalamic Tract<br>Brainstem: Left Occipital Corticopontine Tract<br>Brainstem: Left Parietal Corticopontine Tract<br>Association: Left Inferior Longitudinal Fasciculus<br>Brainstem: Right Parietal Corticopontine Tract<br>Basal Ganglia: Left Posterior Thalamic Radiation<br>Brainstem: Right Medial Lemniscus<br>Association: Left Inferior Fronto-Occipital Fasciculus<br>Basal Ganglia: Left Superior Thalamic Radiation<br>Brainstem: Right Corticospinal Tract<br>Basal Ganglia: Right Superior Corticostriatal Tract<br>Basal Ganglia: Right Posterior Thalamic Radiation | Brainstem: Left Corticospinal Tract<br>Brainstem: Right Corticospinal Tract<br>Brainstem: Right Parietal Corticopontine Tract<br>Brainstem: Left Parietal Corticopontine Tract<br>Brainstem: Left Medial Lemniscus<br>Brainstem: Right Medial Lemniscus<br>Commissure: Corpus Callosum Forceps Major<br>Basal Ganglia: Left Superior Thalamic Radiation<br>Association: Right Extreme Capsule<br>Brainstem: Right Occipital Corticopontine Tract<br>Basal Ganglia: Left Posterior Corticostriatal Tract<br>Basal Ganglia: Right Superior Thalamic Radiation<br>Association: Left Extreme Capsule<br>Commissure: Corpus Callosum Body<br>Basal Ganglia: Left Superior Corticostriatal Tract<br>Association: Left Middle Longitudinal Fasciculus |

| |
|---|
| Brainstem: Left Medial Lemniscus<br>Brainstem: Non-Decussating Dentatorubrothalamic Tract |

*Concussed tracts with a frequency above the control tracts are highlighted in red.*

**Table S13. Full List of Female Concussed and Female Control Tracts from Differential Tractography for QA 0.30**

| Concussed | Control |
|---|---|
| Commissure: Corpus Callosum Forceps Major<br>Commissure: Corpus Callosum Tapetum<br>Association: Left Arcuate Fasciculus<br>Association: Right Inferior Longitudinal Fasciculus<br>Association: Left Parietal Aslant Tract<br>Basal Ganglia: Right Posterior Corticostriatal Tract<br>Commissure: Occipital Anterior Commissure<br>Basal Ganglia: Left Optic Radiation<br>Brainstem: Left Occipital Corticopontine Tract<br>Basal Ganglia: Left Posterior Thalamic Radiation<br>Association: Right Inferior Fronto-Occipital Fasciculus<br>Basal Ganglia: Left Posterior Corticostriatal Tract | Brainstem: Left Parietal Corticopontine Tract<br>Commissure: Corpus Callosum Forceps Major<br>Brainstem: Left Corticospinal Tract<br>Brainstem: Left Medial Lemniscus |

*Concussed tracts with a frequency above the control tracts are highlighted in red. 1/10 (10.0%) of female concussed participants and 1/9 (11.1%) of female control participants had differential tractography results.*

**Table S14. Full List of Female Concussed and Female Control Tracts from Differential Tractography for QA 0.35**

| Concussed | Control |
|---|---|
| Commissure: Corpus Callosum Forceps Major<br>Association: Right Inferior Longitudinal Fasciculus<br>Commissure: Corpus Callosum Tapetum<br>Basal Ganglia: Left Posterior Thalamic Radiation<br>Basal Ganglia: Right Posterior Corticostriatal Tract | N/A |

*Concussed tracts with a frequency above the control tracts are highlighted in red. N/A indicates that no participants in that group had results. 1/10 (10.0%) of female concussed participants and 0/9 (0.0%) of female control participants had differential tractography results.*

**Table S15. Full List of Male Concussed and Male Control Tracts from Differential Tractography for RDI 0.25**

| Concussed | Control |
|---|---|
| Cerebellum: Left Cerebellum | Cerebellum: Right Cerebellum |
| Cerebellum: Right Cerebellum | Cerebellum: Left Cerebellum |
| Cerebellum: Vermis | Cerebellum: Vermis |
| Commissure: Corpus Callosum Forceps Major | Commissure: Corpus Callosum Forceps Major |
| Association: Left Vertical Occipital Fasciculus | Brainstem: Left Medial Lemniscus |
| Association: Left Inferior Fronto-Occipital Fasciculus | Brainstem: Left Medial Forebrain Bundle |
| Commissure: Corpus Callosum Body | Cerebellum: Middle Cerebellar Peduncle |
| Association: Left Inferior Longitudinal Fasciculus | |
| Association: Right Inferior Longitudinal Fasciculus | |
| Association: Right Inferior Fronto-Occipital Fasciculus | |
| Commissure: Occipital Anterior Commissure | |
| Cerebellum: Middle Cerebellar Peduncle | |
| Brainstem: Left Corticospinal Tract | |
| Association: Left Parietal Aslant Tract | |
| Association: Left Arcuate Fasciculus | |
| Brainstem: Left Parietal Corticopontine Tract | |
| Brainstem: Right Corticospinal Tract | |
| Basal Ganglia: Left Superior Thalamic Radiation | |
| Basal Ganglia: Left Optic Radiation | |
| Basal Ganglia: Left Superior Corticostriatal Tract | |
| Brainstem: Non-Decussating Dentatorubrothalamic Tract | |
| Basal Ganglia: Left Posterior Thalamic Radiation | |
| Brainstem: Right Medial Lemniscus | |
| Association: Left Superior Longitudinal Fasciculus | |
| Basal Ganglia: Left Posterior Corticostriatal Tract | |
| Basal Ganglia: Right Posterior Corticostriatal Tract | |
| Commissure: Corpus Callosum Tapetum | |
| Brainstem: Left Medial Lemniscus | |

| |
|---|
| Brainstem: Left Occipital Corticopontine Tract<br>Basal Ganglia: Right Optic Radiation<br>Brainstem: Right Parietal Corticopontine Tract<br>Association: Left Extreme Capsule<br>Association: Left Frontal Aslant Tract<br>Association: Left Superior Longitudinal Fasciculus<br>Basal Ganglia: Right Posterior Thalamic Radiation<br>Association: Right Superior longitudinal Fasiculus Cingulum<br>Basal Ganglia: Right Superior Corticostriatal Tract<br>Basal Ganglia: Right Superior Thalamic Radiation<br>Association: Left Middle Longitudinal Fasciculus<br>Brainstem: Non-Decussating Dentatorubrothalamic Tract<br>Association: Right Vertical Occipital Fasciculus<br>Brainstem: Right Occipital Corticopontine Tract<br>Association: Right Parietal Aslant Tract<br>Brainstem: Right Reticular Tract<br>Brainstem: Right Medial Forebrain Bundle<br>Brainstem: Left Reticular Tract |

*Concussed tracts with a frequency above the control tracts are highlighted in red.*

**Table S16. Full List of Male Concussed and Male Control Tracts from Differential Tractography for RDI 0.30**

| Concussed | Control |
|---|---|
| Cerebellum: Left Cerebellum<br>Commissure: Corpus Callosum Forceps Major<br>Cerebellum: Vermis<br>Association: Right Inferior Longitudinal Fasciculus<br>Association: Left Inferior Fronto-Occipital Fasciculus<br>Association: Left Inferior Longitudinal Fasciculus<br>Basal Ganglia: Left Posterior Thalamic Radiation<br>Cerebellum: Middle Cerebellar Peduncle<br>Association: Left Vertical Occipital Fasciculus<br>Basal Ganglia: Right Posterior Corticostriatal Tract<br>Basal Ganglia: Right Posterior Thalamic Radiation<br>Brainstem: Left Occipital Corticopontine Tract<br>Cerebellum: Right Cerebellum | Cerebellum: Vermis<br>Cerebellum: Right Cerebellum<br>Cerebellum: Left Cerebellum |

*Concussed tracts with a frequency above the control tracts are highlighted in red. 6/14 (42.9%) of male concussed participants and 2/15 (13.3%) of male control participants had differential tractography results.*

**Table S17. Full List of Male Concussed and Male Control Tracts from Differential Tractography for RDI 0.35**

| Concussed | Control |
|---|---|
| Cerebellum: Left Cerebellum<br>Cerebellum: Middle Cerebellar Peduncle<br>Cerebellum: Right Cerebellum | Cerebellum: Right Cerebellum<br>Cerebellum: Left Cerebellum<br>Cerebellum: Vermis |

*Concussed tracts with a frequency above the control tracts are highlighted in red. 3/14 (21.4%) of male concussed participants and 2/15 (13.3%) of male control participants had differential tractography results.*

**Table S18. Full List of Female Concussed and Female Control Tracts from Differential Tractography for RDI 0.25**

| Concussed | Control |
|---|---|
| Cerebellum: Right Cerebellum | Cerebellum: Left Cerebellum |
| Cerebellum: Vermis | Cerebellum: Right Cerebellum |
| Cerebellum: Left Cerebellum | Cerebellum: Middle Cerebellar Peduncle |
| Cerebellum: Middle Cerebellar Peduncle | Cerebellum: Vermis |
| Commissure: Corpus Callosum Forceps Major | Brainstem: Left Corticospinal Tract |
| Association: Left Parietal Aslant Tract | Association: Left Superior Longitudinal Fasciculus |
| Association: Left Vertical Occipital Fasciculus | Brainstem: Right Corticospinal Tract |
| Association: Left Inferior Longitudinal Fasciculus | Basal Ganglia: Left Superior Corticostriatal Tract |
| Association: Left Inferior Fronto-Occipital Fasciculus | Commissure: Corpus Callosum Body |
| Association: Right Inferior Fronto-Occipital Fasciculus | Association: Left Extreme Capsule |
| Association: Right Inferior Longitudinal Fasciculus | Association: Left Inferior Longitudinal Fasciculus |
| Basal Ganglia: Left Posterior Corticostriatal Tract | Brainstem: Left Medial Lemniscus |
| Basal Ganglia: Left Posterior Thalamic Radiation | Brainstem: Left Parietal Corticopontine Tract |
| Association: Left Superior Longitudinal Fasciculus | Association: Left Uncinate Fasciculus |
| Association: Right Vertical Occipital Fasciculus | Basal Ganglia: Right Superior Corticostriatal Tract |
| Association: Left Middle Longitudinal Fasciculus | Association: Left Parietal Aslant Tract |
| Commissure: Occipital Anterior Commissure | Association: Left Arcuate Fasciculus |
| Brainstem: Left Parietal Corticopontine Tract | Brainstem: Right Medial Lemniscus |
| Basal Ganglia: Left Optic Radiation | Basal Ganglia: Left Superior Thalamic Radiation |
| Basal Ganglia: Right Optic Radiation | Brainstem: Right Parietal Corticopontine Tract |
| Commissure: Corpus Callosum Body | Commissure: Temporal Anterior Commissure |
| Commissure: Corpus Callosum Tapetum | Association: Left Middle Longitudinal Fasciculus |
| Association: Left Arcuate Fasciculus | Basal Ganglia: Right Superior Thalamic Radiation |
| Brainstem: Left Occipital Corticopontine Tract | Brainstem: Right Non-Decussating Dentatorubrothalamic Tract |
| Basal Ganglia: Right Posterior Corticostriatal Tract | Association: Left Hippocampus Alveus |
| Basal Ganglia: Left Superior Thalamic Radiation | Association: Right Extreme Capsule |
| Basal Ganglia: Right Posterior Thalamic Radiation | Brainstem: Left Non-Decussating Dentatorubrothalamic Tract |
| | Basal Ganglia: Left Posterior Corticostriatal Tract |
| | Association: Left Superior Longitudinal Fasciculus |
| | Association: Left Frontal Aslant Tract |

| | |
|---|---|
| Association: Left Superior Longitudinal Fasciculus<br>Brainstem: Right Occipital Corticopontine Tract<br>Cerebellum: Right Inferior Cerebellar Peduncle | Brainstem: Left Occiptial Corticopontine Tract<br>Brainstem: Right Frontal Corticopontine Tract<br>Basal Ganglia: Left Posterior Thalamic Radiation<br>Association: Right Superior Longitudinal Fasciculus Cingulum<br>Brainstem: Right Medial Forebrain Bundle<br>Association: Right Frontal Aslant Tract<br>Association: Right Superior Longitudinal Fasciculus<br>Commissure: Corpus Callosum Forceps Major<br>Commissure: Corpus Callosum Tapetum<br>Association: Left Superior Longitudinal Fasciculus Cingulum<br>Brainstem: Right Occipital Corticopontine Tract<br>Commissure: Occipital Anterior Commissure<br>Association: Left Inferior Fronto-Occipital Fasciculus<br>Brainstem: Left Corticobulbar Tract |

*Concussed tracts with a frequency above the control tracts are highlighted in red.*

**Table S19. Full List of Female Concussed and Female Control Tracts from Differential Tractography for RDI 0.30**

| Concussed | Control |
|---|---|
| Cerebellum: Right Cerebellum | Cerebellum: Left Cerebellum |
| Cerebellum: Left Cerebellum | Cerebellum: Middle Cerebellar Peduncle |
| Commissure: Corpus Callosum Forceps Major | Association: Left Superior Longitudinal Fasciculus |
| Association: Left Vertical Occipital Fasciculus | Brainstem: Left Corticospinal Tract |
| Association: Left Inferior Fronto-Occipital Fasciculus | Association: Left Extreme Capsule |
| Association: Left Inferior Longitudinal Fasciculus | Brainstem: Left Parietal Corticopontine Tract |
| Association: Left Parietal Aslant Tract | Commissure: Corpus Callosum Body |
| Basal Ganglia: Left Posterior Corticostriatal Tract | Brainstem: Left Medial Lemniscus |
| Association: Right Inferior Fronto-Occipital Fasciculus | Association: Left_Middle Longitudinal Fasciculus |
| Basal Ganglia: Left Optic Radiation | Basal Ganglia: Left Superior Thalamic Radiation |
| Brainstem: Left Parietal Corticopontine Tract | Brainstem: Right Corticospinal Tract |
| Basal Ganglia: Left Posterior Thalamic Radiation | Basal Ganglia: Left Superior Corticostriatal Tract |
| Commissure: Occipital Anterior Commissure | Brainstem: Right Parietal Corticopontine Tract |
| Association: Left Superior Longitudinal Fasciculus | Brainstem: Right Medial Lemniscus |
| Commissure: Corpus Callosum Body | Basal Ganglia: Left Posterior Thalamic Radiation |
| Association: Left Middle Longitudinal Fasciculus | Basal Ganglia: Left Posterior Corticostriatal Tract |
| Association: Right Inferior Longitudinal Fasciculus | Association: Right Extreme Capsule |
| Brainstem: Left Occipital Corticopontine Tract | Basal Ganglia: Right Superior Corticostriatal Tract |
| Association: Right Vertical Occipital Fasciculus | Brainstem: Left Non-Decussating Dentatorubrothalamic Tract |
| Basal Ganglia: Right Posterior Corticostriatal Tract | Brainstem: Right Non-Decussating Dentatorubrothalamic Tract |
| Basal Ganglia: Right Optic Radiation | |
| Association: Left Arcuate Fasciculus | |
| Cerebellum: Vermis | |
| Cerebellum: Right Inferior Cerebellar Peduncle | |

*Concussed tracts with a frequency above the control tracts are highlighted in red. 3/10 (30.0%) of female concussed participants and 3/9 (33.3%) of female control participants had differential tractography results.*

**Table S20. Full List of Female Concussed and Female Control Tracts from Differential Tractography for RDI 0.35**

| Concussed | Control |
|---|---|
| Cerebellum: Right Cerebellum<br>Association: Left Vertical Occipital Fasciculus<br>Association: Left Inferior Fronto-Occipital Fasciculus<br>Commissure: Corpus Callosum Forceps Major<br>Association: Left Inferior Longitudinal Fasciculus<br>Cerebellum: Left Cerebellum<br>Basal Ganglia: Left Posterior Corticostriatal Tract<br>Basal Ganglia: Left Optic Radiation<br>Basal Ganglia: Left Posterior Thalamic Radiation<br>Brainstem: Left Occipital Corticopontine Tract<br>Commissure: Occipital Anterior Commissure | Cerebellum: Left Cerebellum<br>Cerebellum: Middle Cerebellar Peduncle |

*Concussed tracts with a frequency above the control tracts are highlighted in red. 2/10 (20.0%) of female concussed participants and 2/9 (22.2%) of female control participants had differential tractography results.*

**Table S21. FDR Values for QA and RDI Increases at 25%, 30%, and 35% Thresholds**

| | 25% | 30% | 35% |
|---|---|---|---|
| **QA** | 0.363 | 0.004 | - |
| **RDI** | 13.805 | 31.533 | 123.556 |

**Table S22. Full List of Concussed and Control Tracts from Differential Tractography for Increased QA 0.30**

| Concussed | Control |
|---|---|
| Commissure: Corpus Callosum Forceps Minor | N/A |

*Concussed tracts with a frequency above the control tracts are highlighted in red. N/A indicates that no participants in that group had results. 1/24 (4.2%) of concussed participants and 0/24 (0.00%) of control participants had differential tractography results.*
